# Ionomycin Ameliorates Hypophosphatasia *via* Rescuing Alkaline Phosphatase Deficiency-mediated L-type Ca^2+^ Channel Internalization in Mesenchymal Stem Cells

**DOI:** 10.1101/545418

**Authors:** Bei Li, Xiaoning He, Zhiwei Dong, Kun Xuan, Wei Sun, Li Gao, Shiyu Liu, Wenjia Liu, Chenghu Hu, Yimin Zhao, Songtao Shi, Yan Jin

**Affiliations:** State Key Laboratory of Military Stomatology&National Clinical Research Center for Oral Diseases&Shaanxi International Joint Research Center for Oral Diseases, Center for Tissue Engineering, School of Stomatology, The Fourth Military Medical University, Xi’an, Shaanxi 710032, China; Xi’an Institute of Tissue Engineering and Regenerative Medicine, Xi’an, China; Department of Anatomy and Cell Biology, University of Pennsylvania, School of Dental Medicine, Philadelphia, PA, USA; Department of Oral and Maxillofacial surgery, General Hospital of Shenyang Military Area Command, Shenyang, Liaoning, China; Department of Pediatric Dentistry, School of Stomatology, Fourth Military Medical University, Xi’an, China; Institute for Biomedical Sciences of Pain, Tangdu Hospital, Fourth Military Medical University, Xi’an, China.

**Author notes:** Corresponding Author: Professor Yan Jin, MS-State Key Laboratory, School of Stomatology, Fourth Military Medical University, Xi’an, Shaanxi 710032, China, or Professor Songtao Shi, South China Center of Craniofacial Stem Cell Research, Guanghua School of Stomatology, Sun Yat-sen University. 74 Zhongshan 2Rd, Guangzhou, Guangdong, China,. These authors contributed equally to this work (Bei Li, Xiaoning He, Zhiwei Dong). South China Center of Craniofacial Stem Cell Research, Guanghua School of Stomatology, Sun Yat-sen University. 74 Zhongshan 2Rd, Guangzhou, Guangdong, China.

## Abstract

Loss-of-function mutations in ALPL result in hypophosphatasia (HPP), an inborn error of metabolism that causes skeletal mineralization defect. In adults, main clinical involvement includes early loss of primary or secondary teeth, osteoporosis, bone pain, chondrocalcinosis, and fractures. However, guidelines for the treatment of adults with HPP are not available. Here, we show that ALPL deficiency caused reduction of intracellular Ca^2+^ influx resulting in osteoporotic phenotype due to downregulated osteogenic differentiation and upregulated adipogenic differentiation in both human and mouse BMSCs. To elevate intracellular level of calcium in bone marrow mesenchymal stem cells (BMSCs) by ionomycin treatment rescues the osteoporotic phenotype in *alpl*^*+/-*^ mice and BMSC-specific (*Prrx1-alpl*^*-/-*^) conditional alpl knockout mice. Mechanistically, ALPL is required to maintain intracellular Ca^2+^ influx by regulating L-type Ca^2+^ channel trafficking *via* binding to the α2δ subunits, which regulates the internalization of L-type Ca^2+^ channel. Decreased Ca^2+^ flux inactivates Akt/GSK3β/β-catenin signaling pathway that regulates lineage differentiation of BMSCs. This study identifies a previous unknown role of ectoenzyme ALPL in maintenance of calcium channel trafficking to keep stem cell lineage differentiation and bone homeostasis. Accelerating Ca^2+^ flux through L-type Ca^2+^ channel by ionomycin treatment may be a promising therapeutic approach for adult HPP patients.

**One Sentence Summary:** ALP regulates internalization of L-Type Ca^2+^ Channel of BMSCs in Hypophosphatasia.

## Introduction

A loss of function mutation in the liver/bone/kidney alkaline phosphatase (ALPL) gene results in life-threatening diseases of hypophosphatasia (HPP) during early developmental periods, featuring hypomineralization of skeleton and teeth (Millan & Whyte, 2016; Whyte, 2016). Adult HPP patients showed early loss of primary or secondary teeth, osteoporosis, bone pain, chondrocalcinosis, and fractures. Our previous study found age-related bone mass loss and marrow fat gain in heterozygous *Alpl*^*+/-*^ mice (Liu et al, 2018). Bone marrow mesenchymal stem cells (BMSCs) are multipotent cells capable of differentiating into various cell lineages including osteoblasts and adipoytes. With age, BMSCs are more inclined to undergo differentiation into adipocytes rather than osteoblast, resulting in an increased number of adipocytes and a decresed number of osteoblasts, causing osteoporosis (Li et al, 2018). Our previous study also showed that ALPL governed the osteo-adipogenic balance of BMSCs and prevent cell senescence (Liu et al, 2018). Alkaline phosphatase is known as a ubiquitous plasma membrane-bound enzyme (ectoenzyme), which hydrolyzes several different molecules at physiological (neutral) and alkaline pH, including inorganic pyrophosphate (Russell, 1965), pyridoxal-5-phosphate (the active form of vitamin B6) (Whyte et al, 1985) and nucleotides (Ciancaglini et al, 2010; Scheibe et al, 2000; Zimmermann, 2006). However, the detail mechanism of ALPL causing this age-related osteoporosis is largely unknown.

In severely affected infantile HPP, hypercalcemia and hypercalciuria were often reported as symptoms (Barcia et al, 1997; Belachew et al, 2013). However, it is still not clear why calcium metabolism abnormalities are induced by ALPL mutation, as it plays an important role in generating the inorganic phosphate. Meanwhile, whether the aberrant calcium metabolism involved in age-related osteoporosis of heterozygous *Alpl*^*+/-*^ mice is also unclear. It is well accepted that calcium metabolism abnormalities are closely related with calcium channel on the cell surface. Calcium influx is controlled by voltage-gated Ca^2+^ channels (VGCCs), or agonist-dependent and voltage-independent Ca^2+^ entry pathways, which are called ‘store-operated’ Ca^2+^ channels (SOCs). Changes in intracellular Ca^2+^ concentration ([Ca^2+^]_i_) play an essential role in regulating motility, apoptosis, differentiation and many other cellular processes (Berridge et al, 2000). Aberrant intracellular [Ca^2+^]_i_ leads to loss of Ca^2+^ homeostasis, which causes abnormal calcium metabolism and bone disorder (Cui et al, 2012; Hoenderop et al, 2003). Several types of Ca^2+^ channels are reported to regulate intracellular Ca^2+^ homeostasis in BMSCs and osteoblasts to affect bone repair (Barradas et al, 2012; Jung et al, 2015; Wen et al, 2012). Thus, the regulation of Ca^2+^ channels at the membrane plays a central role in BMSC function and bone related diseases. However, whether ALPL modulates Ca^2+^ channels to maintain Ca^2+^ homeostasis in BMSCs is unknown.

Currently, specific medical treatment options for HPP are limited to bone-targeted enzyme replacement therapy (asfotase, Strensiq, Alexion), approved for pediatric-onset HPP (Whyte et al, 2012; Whyte et al, 2016). At this time, there are no guidelines for selecting adult patients for treatment, for evaluating the results of treatment, or determining the optimal duration of treatment. In this study, we use human and mouse model to demonstrate that ALPL is required to maintain intracellular Ca^2+^ influx by regulating L-type Ca^2+^ channel trafficking *via* binding to the α2δ subunits, which regulates the internalization of L-type Ca^2+^ channel. This decreased Ca^2+^ flux downregulates Akt/GSK3β-mediated Wnt/β-catenin signaling in BMSCs, leading to age-related osteoporotic phenotype. Moreover, we found that raising the intracellular level of calcium in BMSCs by ionomycin treatment rescues the osteoporotic phenotype in *alpl*^*+/-*^ and BMSC-specific (*Prrx1-alpl*^*-/-*^) conditional *alpl* knockout mice, as well as stem cell function of BMSCs from HPP patients, suggesting a new strategy for HPP therapy.

## Results

### ALPL deficiency caused decreased membrane expression of L-type Ca^2+^ channel in BMSCs

As severe ALPL deficiency patients develop hypercalcemia (Barcia et al, 1997; Belachew et al, 2013), we examined the plasma calcium level in *alpl*^*+/-*^ mice and found a marked increase in the level of plasma calcium (**Fig S1a**). To explore whether ALPL deficiency contributes to abnormal calcium metabolism in BMSCs, we isolated BMSCs from *alpl*^*+/-*^ mice (**Fig S1b-e**) and examined the cytosolic Ca^2+^. We found that lentivirus-overexpressing ALPL in *alpl*^*+/-*^ BMSCs is able to increase the cytosolic Ca^2+^ of *alpl*^*+/-*^ BMSCs (**Fig S2a**). Ca^2+^ entry across the plasma membrane occurs *via* two distinct pathways, SOCs and VGCCs. To test which type Ca^2+^ channels might be affected in the absence of ALPL, WT and *alpl*^*+/-*^ BMSCs were cultured with 10 nM thapsigargin (TG), a noncompetitive inhibitor capable of raising cytosolic Ca^2+^ concentration *via* blocking the ability of the cell to pump Ca^2+^ into the sarcoplasmic and endoplasmic reticula, to activate plasma membrane Ca^2+^ channels. Evidently, no significant difference of TG-induced intracellular Ca^2+^ influx was detected in WT, *alpl*^*+/-*^ and shALP BMSCs (**Fig S2b, c**). These data suggest that ALPL deficiency affects the VGCC function in BMSCs. The contribution of ALPL to the VGCCs in BMSCs was determined after membrane depolarization using 30 mM KCl. Intracellular Ca^2+^ imaging analysis showed that KCl-induced Ca^2+^ influx was significantly decreased in culture-expanded *alpl*^*+/-*^ and shALP BMSCs compared to WT BMSCs (**Fig 1a**). Moreover, intracellular Ca^2+^ imaging analysis showed that KCl-induced Ca^2+^ influx was not changed in culture-expanded WT, *alpl*^*+/-*^ and shALP BMSCs when treated with 10 mM EGTA (**Fig 1b**), suggesting that ALPL regulates Ca^2+^ elevation mainly due to Ca^2+^ influx with a limited contribution from intracellular Ca^2+^ storage. Overexpression of ALPL in *alpl*^*+/-*^ BMSCs rescued KCl-induced Ca^2+^ influx (**Fig 1c**).

**Figure 1.**
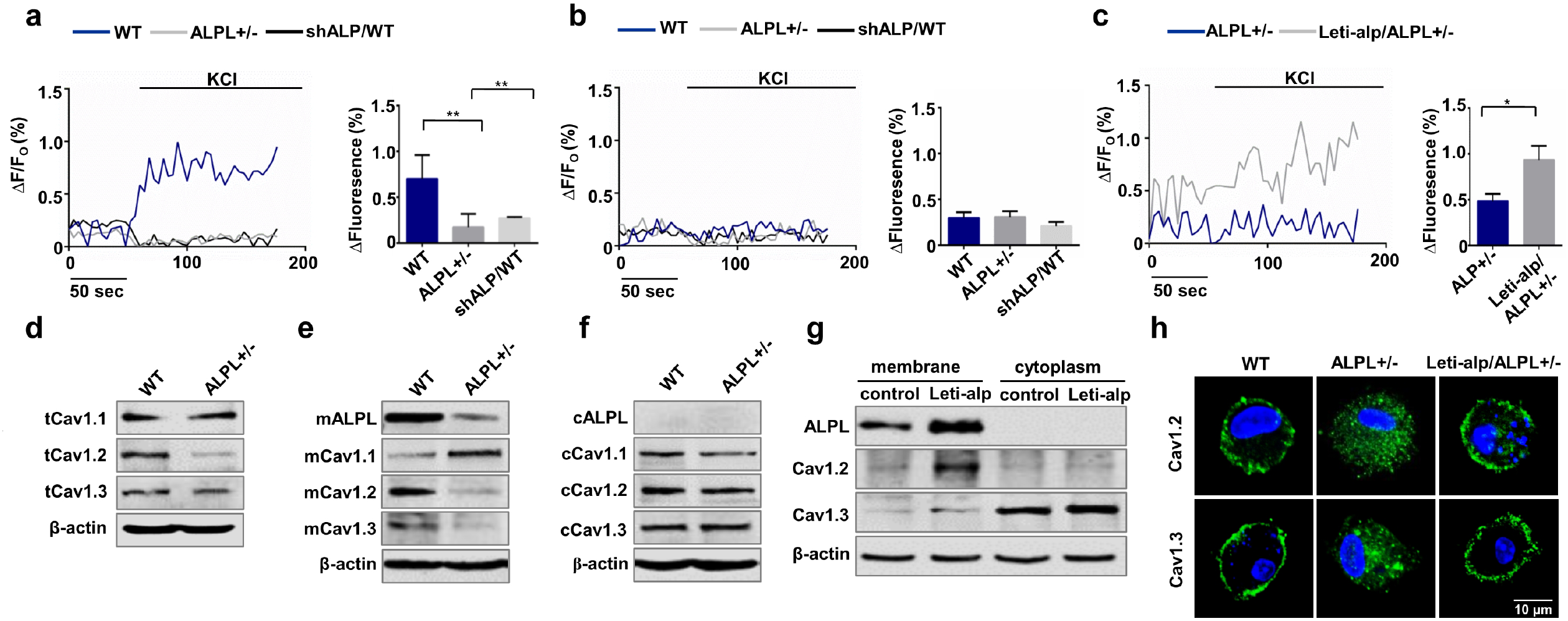
ALPL deficiency caused decreased membrane expression of L-type Ca^2+^ channel in BMSCs. (a) Ca^2+^ imaging showed decreased Ca^2+^ influx of cultured *alpl*^*+/-*^ BMSCs and WT BMSCs transfected with shALP (shALP/WT) stimulated with 30 mM KCl for 3 min (n=10). (b) No KCl-induced Ca^2+^ influx was detected in cultured WT, *alpl*^*+/-*^ and shALP/WT BMSCs treated by 10 mM EGTA for 3 min (n=10). (c) Lentivirus-overexpressing ALPL treated *alpl*^*+/-*^ (Leti-alp/*alpl*^*+/-*^) BMSCs showed an elevated Ca^2+^ influx stimulated with 30 mM KCl for 3 min (n=10). (d, e, f) The expression of Ca_V_1.1, Ca_V_1.2 and Ca_V_1.3 was assessed. *alpl*^*+/-*^ BMSCs showed downregulation of total cell expression (d) and membrane expression of Ca_V_1.2 and Ca_V_1.3 (e), and no change of cytoplasm expression of Ca_V_1.2 and Ca_V_1.3 (f). (g) Leti-alp/*alpl*^*+/-*^ BMSCs showed an elevated membrane expression of ALP, Ca_V_1.2 and Ca_V_1.3. (h) Representative images of confocal laser scanning microscope showed membrane location of Ca_V_1.2 and Ca_V_1.3 in WT and Leti-alp/*alpl*^*+/-*^ BMSCs. Scale bar, 10 μm. Representative results were from three independent experiments. Error bars represent the s.d. from the mean values. *P < 0.05. **P < 0.01. *** P < 0.001.

The decrease in Ca^2+^ currents in *alpl*^*+/-*^ BMSCs could arise from loss of channels at the membrane. The VGCCs comprise 10 subsets, Ca_V_1.1, Ca_V_1.2, Ca_V_1.3, Ca_V_1.4 (L-type), Ca_V_2.1 (P/Q-type), Ca_V_2.2 (N-type), Ca_V_2.3 (R-type), Ca_V_3.1, Ca_V_3.2 and Ca_V_3.3 (T-type) encoded by cacna1s, cacna1c, cacna1d, cacna1f, cacna1a, cacna1b, cacna1e, cacna1g, cacna1h and cacna1i, respectively (Catterall, 2000; Tsien et al, 1988). To identify which subunits of the VGCCs were regulated by ALPL, we measured total protein expression of Ca_V_1.1, Ca_V_1.2, Ca_V_1.3, Ca_V_2.1, Ca_V_2.2, Ca_V_2.3, Ca_V_3.1, Ca_V_3.2 and Ca_V_3.3, as Ca_V_1.4 seems so far to be restricted to the retina. The results showed that total protein expression of Ca_V_1.2 and Ca_V_1.3 were decreased significantly in *alpl*^*+/-*^ BMSCs compared to WT BMSCs (**Fig 1d**). Total protein expression of Ca_V_1.1, Ca_V_2.2, Ca_V_2.3 and Ca_V_3.3 were not changed (**Fig 1d and Fig S2d**). However, total and membrane expression of Ca_V_2.1, Ca_V_3.1 and Ca_V_3.2 were increased in *alpl*^*+/-*^ BMSCs compared to WT BMSCs (**Fig S2d, e**). The expression of Ca_V_2.1, Ca_V_3.1 and Ca_V_3.2 in cytoplasm was not changed significantly in *alpl*^*+/-*^ BMSCs compared to WT BMSCs (**Fig S2f**). Membrane expression of Ca_V_1.1 was increased but the expression in cytoplasm was decreased (**Fig 1e, f**). Considering the decreased Ca^2+^ influx in *alpl*^*+/-*^ BMSCs, we focused on Ca_V_1.2 and Ca_V_1.3. Membrane expression of calcium channel affects the calcium influx, we compared the expression of Ca_V_1.2 and Ca_V_1.3 on membrane and in cytoplasm. The results showed that the expression of Ca_V_1.2 and Ca_V_1.3 on membrane was decreased in *alpl*^*+/-*^ BMSCs compared to WT BMSCs (**Fig 1e**). However, the expression of Ca_V_1.2 and Ca_V_1.3 in cytoplasm was not changed significantly in *alpl*^*+/-*^ BMSCs compared to WT BMSCs (**Fig 1f**). Overexpression of ALPL increased the expression of Ca_V_1.2 and Ca_V_1.3 on membrane as assayed by western blot (**Fig 1g**). We also investigated the membrane localization of Ca_V_1.2 and Ca_V_1.3 in WT and *alpl*^*+/-*^ BMSCs by confocal laser scanning microscope. The results showed that Ca_V_1.2 (FITC labeled) and Ca_V_1.3 (FITC labeled) were localized on the membrane of WT BMSCs (**Fig 1h**). However, Ca_V_1.2 and Ca_V_1.3 were absent on the membrane in *alpl*^*+/-*^ BMSCs. Ca_V_1.2 and Ca_V_1.3 were localized on cell membrane after overexpression of ALPL in *alpl*^*+/-*^ BMSCs (**Fig 1h**). These results suggest that ALPL modulates the member expression of L-type Ca^2+^ channel, especially Ca_V_1.2 and Ca_V_1.3.

### ALPL maintained MSC osteogenic/adipogenic lineage differentiation *via* L-type Ca^2+^ channel

We knocked down ALPL in BMSCs by siRNA (**Fig S3a**) and confirmed that ALPL deficiency decreased the osteogenic differentiation and increased the adipocenic differentiation of BMSCs (**Fig S3b-e**). To address whether the defective osteogenic/adipogenic linage differentiation of BMSCs with ALPL deficiency owing to the abnormal membrane expression of VGCCs, especially Ca_V_1.2 and Ca_V_1.3, we overexpressed Ca_V_1.2 and Ca_V_1.3 in *alpl*^*+/-*^ BMSCs (oeCa_V_1.2 or oeCa_V_1.3), and western blot data showed that Ca_V_1.2 and Ca_V_1.3 were increased on membrane of *alpl*^*+/-*^ BMSCs after Ca_V_1.2 or Ca_V_1.3 overexpression (oeCa_V_1.2 or oeCa_V_1.3) (**Fig 2a**). KCl-induced Ca^2+^ influx was elevated after overexpression of Ca_V_1.2 or Ca_V_1.3 in *alpl*^*+/-*^ BMSCs (**Fig 2b**). We also observed the membrane localization of Ca_V_1.2 and Ca_V_1.3 in *alpl*^*+/-*^ BMSCs after overexpressing Ca_V_1.2 or Ca_V_1.3 by confocal laser scanning microscope (**Fig 2c**). We found that overexpression of Ca_V_1.2 or Ca_V_1.3 in *alpl*^*+/-*^ BMSCs (oeCa_V_1.2 or oeCa_V_1.3) rescued decreased osteogenic differentiation of BMSCs, as evidenced by increased mineralized nodule formation and expression of the osteogenic markers RUNX2 and Sp7 (**Fig. 2d, e**). In contrast, overexpression of Ca_V_1.2 or Ca_V_1.3 (oeCa_V_1.2 or oeCa_V_1.3) decreased adipogenic differentiation of BMSCs, as assessed by Oil red O staining to show decreased numbers of adipocytes and western blot to show downregulation of the adipogenic regulators PPARγ2 and LPL under the adipogenic culture conditions (**Fig. 2f, g**). However, knockdown of Ca_V_1.2 or Ca_V_1.3 (siCa_V_1.2 or siCa_V_1.3) in WT BMSCs caused decreased osteogenic and increased adipogenic differentiation (**Fig. 2h-k**). These data indicate that ALPL regulates osteogenic and adipogenic lineage differentiation through Ca_V_1.2- and Ca_V_1.3-mediated calcium influx.

**Figure 2.**
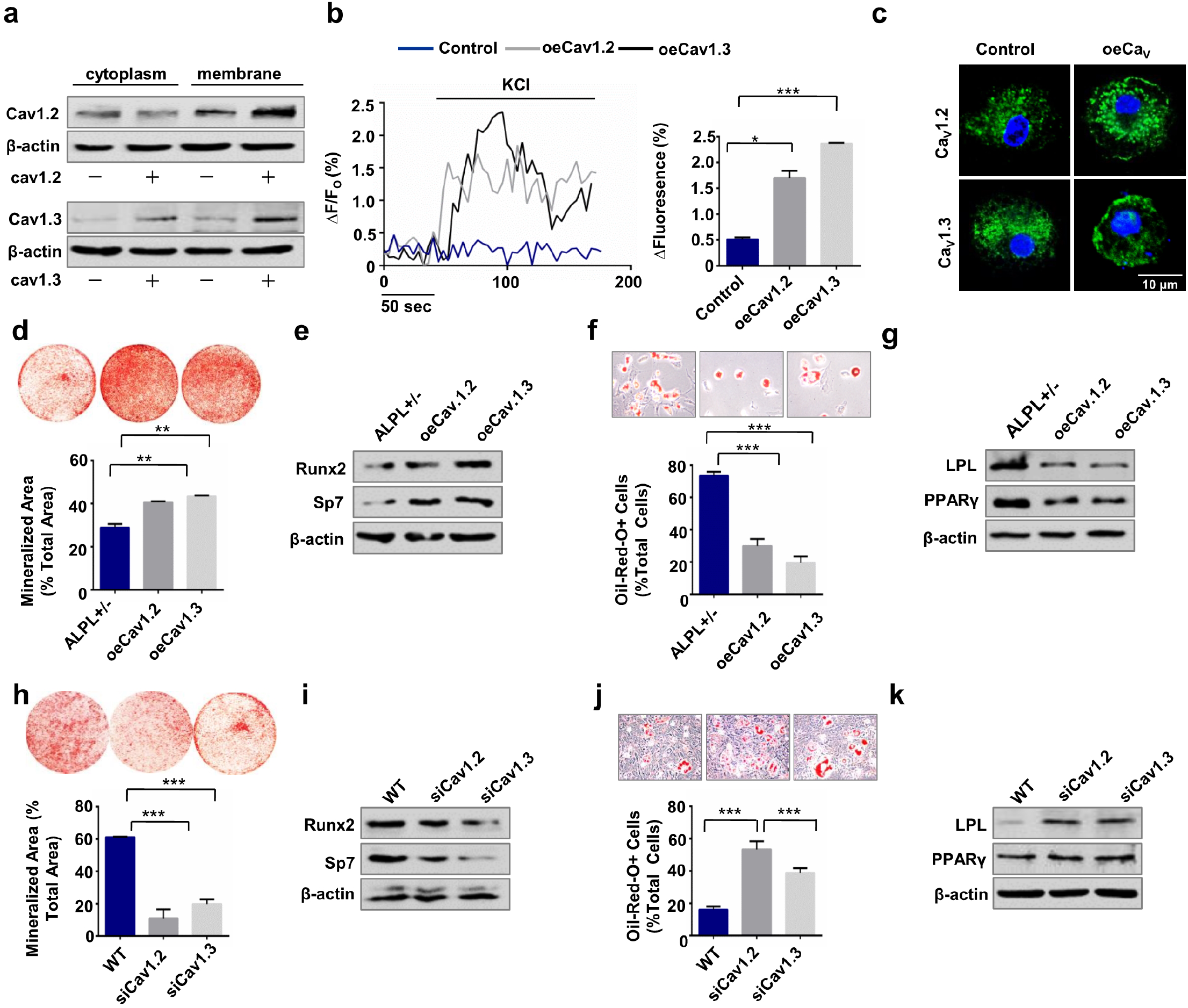
ALPL maintained MSC osteogenic/adipogenic lineage differentiation via L-type Ca^2+^ channel. (a) Plasmid-overexpressed Ca_V_1.2 and Ca_V_1.3 of *alpl*^*+/-*^ BMSCs showed upregulated membrane expression of both channels by western blot analysis. (b) Ca^2+^ imaging showed elevated Ca^2+^ influx of cultured *alpl*^*+/-*^ BMSCs transfected with oeCa_V_1.2 or oeCa_V_1.3 stimulated with 30 mM KCl for 3 min (n=10). (c) Representative images of confocal laser scanning microscope showed membrane location of Ca_V_1.2 and Ca_V_1.3 in *alpl*^*+/-*^ BMSCs transfected with oeCa_V_1.2 or oeCa_V_1.3. Scale bar, 10 μm. (d) Alizarin red staining showed that *alpl*^*+/-*^ BMSCs transfected with siCa_V_1.2 or siCa_V_1.3 had a decreased capacity to form mineralized nodules when cultured under osteo-inductive conditions. (e) Western blot analysis showed that BMSCs transfected with siCa_V_1.2 or siCa_V_1.3 expressed decreased levels of the osteogenic related proteins RUNX2 and Sp7. β-actin was used as a protein loading control. (f, g) BMSCs transfected with siCa_V_1.2 or siCa_V_1.3 showed an increased number of Oil red O-positive adipocytes when cultured under adipo-inductive conditions (f) and upregulation of the adipogenic related proteins PPARγ2 and LPL, as assessed by western blot (g). (h, i) Alizarin red staining showed that *alpl*^*+/-*^ BMSCs transfected with oeCa_V_1.2 or oeCa_V_1.3 had an increased capacity to form mineralized nodules when cultured under osteo-inductive conditions (h) and upregulation of the osteogenic related proteins RUNX2 and Sp7 (i). (j, k) oeCa_V_1.2 or oeCa_V_1.3 treated *alpl*^*+/-*^BMSCs showed a decreased number of Oil red O-positive adipocytes when cultured under adipo-inductive conditions (j) and downregulation of the adipogenic related proteins PPARγ2 and LPL, as assessed by western blot (k). Representative results were from three independent experiments. Error bars represent the s.d. from the mean values. *P < 0.05. **P < 0.01. *** P < 0.001.

### ALPL deficiency promoted the internalization of L-type Ca^2+^ channel in BMSCs

To determine whether lack of ALPL leads to internalization, which caused decreased membrane expression of L-type Ca^2+^ channel in BMSCs, we disrupted endocytosis by expressing a dominant-negative mutant of dynamin 1 (DN-Dyn1), a GTPase required for the formation of endocytic vesicles from the plasma membrane (Praefcke & McMahon, 2004). Expression of DN-Dyn1 in *alpl*^*+/-*^ BMSCs prevented the loss of Ca_V_1.2 and Ca_V_1.3 on the cell surface (**Fig 3a**), providing evidence that lack of ALPL causes internalization of the channels. Western blot analysis showed that expression levels of membrane Ca_V_1.2 and Ca_V_1.3 in *alpl*^*+/-*^ BMSCs were increased after DN-Dyn1 transfection. However, the expression levels of cytoplasm Ca_V_1.2 and Ca_V_1.3 in *alpl*^*+/-*^ BMSCs were not significantly changed after DN-Dyn1 transfection (**Fig 3b**). Expression of DN-Dyn1 also prevented the decrease of KCl-induced Ca^2+^ influx in *alpl*^*+/-*^ BMSCs (**Fig 3c**).

**Figure 3.**
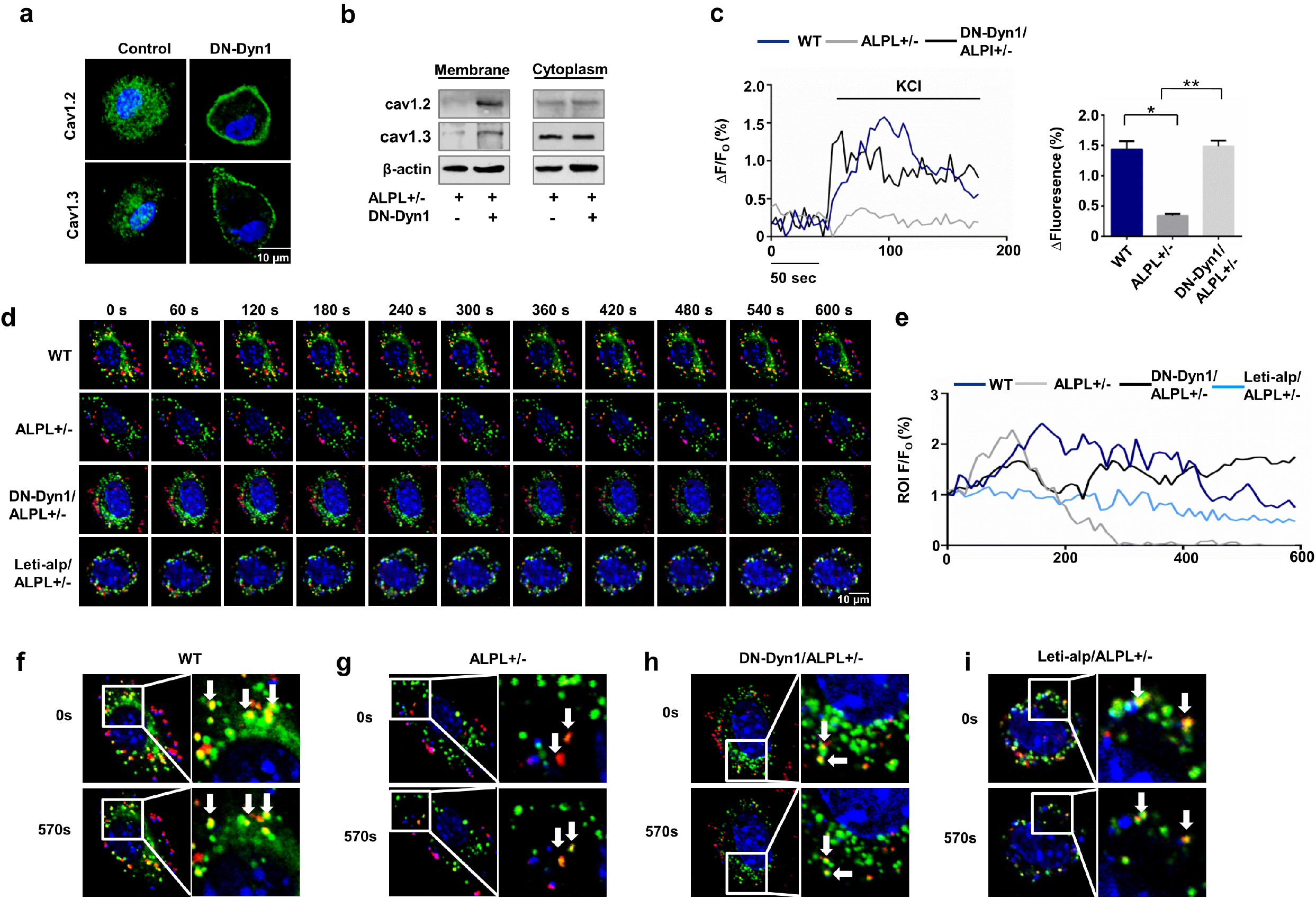
ALPL deficiency promoted the internalization of L-type Ca^2+^ channel in BMSCs. (a) Representative images of confocal laser scanning microscope showed membrane location of Ca_V_1.2 and Ca_V_1.3 in *alpl*^*+/-*^ BMSCs transfected with DN-Dyn1. Scale bar, 10 μm. (b) *alpl*^*+/-*^ BMSCs transfected with DN-Dyn1 showed upregulation of membrane expression of Ca_V_1.2 and Ca_V_1.3 and almost no change of cytoplasm expression of Ca_V_1.2 and Ca_V_1.3, as assessed by western blot. β-actin was used as a protein loading control. (c) Ca^2+^ imaging showed elevated Ca^2+^ influx of cultured *alpl*^*+/-*^ BMSCs transfected with DN-Dyn1 when stimulated with 30 mM KCl for 3 min (n=10). (d, e) 10 min time-lapse confocal laser scanning microscopy images of WT, *alpl*^*+/-*^, DN-Dyn1/*alpl*^*+/-*^ and Leti-alp/*alpl*^*+/-*^ BMSCs containing DsRed-Ca_V_1.2. Scale bar, 10 μm (d) *alpl*^*+/-*^ BMSCs showed a pronounced decrease of DsRed-Ca_V_1.2 on the membrane (Dio labeled ROI, n=10) (e). (f-i) Representative images showed co-localization of DsRed-Ca_V_1.2 with Dio labeled cell membrane of BMSCs. WT, DN-Dyn1/*alpl*^*+/-*^ and Leti-alp/*alpl*^*+/-*^ BMSCs had co-localization region at 0s and 570s (f, h, i). However, *alpl*^*+/-*^ BMSCs showed no co-localization at 0s and 570s (g). Representative results were from three independent experiments. Error bars represent the s.d. from the mean values.*P<0.05, ** P < 0.01.

We next measured the time course of ALPL-dependent L-type Ca^2+^ channel internalization. To study this process in live BMSCs, we used Dio to label the cell membrane (FITC labeled) and constructed a plasmid to express Ca_V_1.2 (DsRed-Cav1.2) and transfect the cells. We recorded co-localization region as region of interest(ROI) to record the time-course change of intensity of red fluorescence. DsRed-Ca_V_1.2 signal in *alpl*^*+/-*^ BMSCs declined after 10 min, reflecting the decreased membrane expression of L-type Ca^2+^ channel (**Fig 3d, e**). However, DsRed-Ca_V_1.2 signal was not changed significantly after 10 min in WT, DN-Dyn1-transfected *alpl*^*+/-*^ BMSCs and ALPL-overexpressed *alpl*^*+/-*^ BMSCs (**Fig 3d, e**). We also selected the images of 0 s and 570 s to show the co-localization of DsRed-Ca_V_1.2 with cell membrane of BMSCs. Almost no co-localization region was found in *alpl*^*+/-*^ BMSCs (**Fig 3f-i**), which suggested ALPL deficiency promoted the internalization of L-type Ca^2+^ channel. To know whether expression of DN-Dyn1 rescues the differentiation in ALPL deficient BMSCs, we performed osteogenic and adipogenic induction after the transfection. Expression of DN-Dyn1 increased osteogenic differentiation of *alpl*^*+/-*^ BMSCs, as evidenced by increased mineralized nodule formation and expression of the osteogenic markers RUNX2 and Sp7 (**Fig s4a, b)**. In contrast, expression of DN-Dyn1 decreased adipogenic differentiation of *alpl*^*+/-*^ BMSCs, as assessed by Oil red O staining to show decreased numbers of adipocytes and western blot to show downregulation of the adipogenic regulators PPARγ2 and LPL under the adipogenic culture conditions (**Fig s4c, d)**.

### ALPL deficiency promoted the internalization of L-type Ca^2+^ channel *via* binding to α2δ subunits

Given that alkaline phosphatase has been reported to hydrolyze inorganic pyrophosphate (PPi) and adenosine triphosphate (ATP), we compared the member expression of Ca_V_1.2 and Ca_V_1.3 in BMSCs and BMSCs treated by PPi or ATP. However, exogenous PPi or ATP barely changed the memberane expression of Ca_V_1.2 and Ca_V_1.3 in BMSCs (**Fig S5a**). To further explore the molecular mechanism of ALPL-regulated internalization of Ca_V_1.2 and Ca_V_1.3, we measured the membrane localization of ALPL and calcium channel. We found that Ca_V_1.2 and Ca_V_1.3 were overlapped with ALPL in BMSCs (**Fig 4a, upper panel**), suggesting the binding of ALPL and L-type Ca^2+^ channel. However, no membrane location of Ca_V_1.2 and Ca_V_1.3 were found in *alpl*^*+/-*^ BMSCs (**Fig 4a, lower panel**). Several co-localization regions of ALPL and Ca_V_1.2 and Ca_V_1.3 were found in cytoplasm of *alpl*^*+/-*^ BMSCs (**Fig 4a, lower panel**), indicating that ALPL may bind L-type Ca^2+^ channel in cytoplasm. We also used immunoprecipitation to confirm the binding of ALPL and L-type Ca^2+^ channel. Immunoprecipitation using a control antibody did not isolate either protein, but an immunoprecipitation with anti-ALPL resulted in a coimmunoprecipitation with Ca_V_1.2 or Ca_V_1.3 (**Fig 4b**). In addition, we found anti-Ca_V_1.2 or anti-Ca_V_1.3 enriched ALPL (**Fig 4c**), suggesting that ALPL directly binds with Ca_V_1.2 and Ca_V_1.3 in BMSCs.

**Figure 4.**
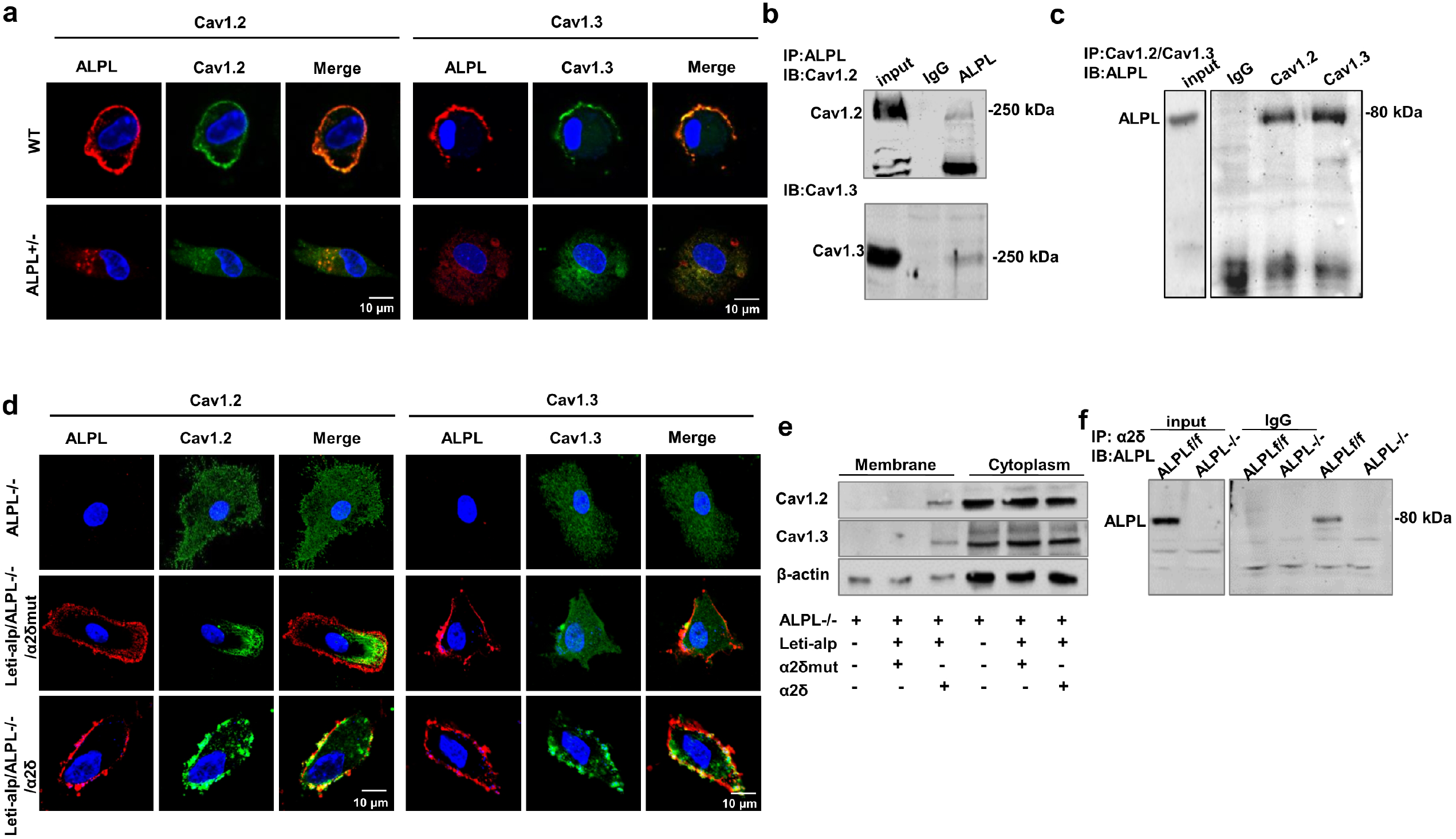
ALPL deficiency promoted the internalization of L-type Ca^2+^ channel *via* binding to α2δ subunits. (a) Representative images of confocal laser scanning microscope showed membrane co-localization region of ALPL (DsRed) and Ca_V_1.2 (FITC-labeled) or Ca_V_1.3 (FITC-labeled) in WT BMSCs. No membrane co-localization region was found in *alpl*^*+/-*^ BMSCs. Scale bar, 10 μm. (b) ALPL immunoprecipitated Ca_V_1.2 and Ca_V_1.3. Left lane showed the expression of Ca_V_1.2 and Ca_V_1.3, and right lane showed the levels of Ca_V_1.2 and Ca_V_1.3 following immunoprecipitation with an anti-ALPL antibody. (c) Ca_V_1.2 and Ca_V_1.3 immunoprecipitated ALPL. Left panel showed the expression of ALPL and right panel showed the level of ALPL following immunoprecipitation with anti-Ca_V_1.2 or anti-Ca_V_1.3 antibody. (d) Representative images of confocal laser scanning microscope showed membrane co-localization region of ALPL (Cy3-labeled) and Ca_V_1.2 (FITC-labeled) or Ca_V_1.3 (FITC-labeled) in *alpl*^*-/-*^ BMSCs overexpressed with ALPL and α2δ subunit. No membrane co-localization region was found in *alpl*^*-/-*^ BMSCs and *alpl*^*-/-*^ BMSCs overexpressed with ALPL and mutant α2δ subunit. Scale bar, 10 μm. (e) Western blot analysis showed membrane expression of Ca_V_1.2 or Ca_V_1.3 in *alpl*^*-/-*^ BMSCs overexpressed with ALPL and α2δ subunit. No membrane expression of Ca_V_1.2 or Ca_V_1.3 was found in *alpl*^*-/-*^ BMSCs and *alpl*^*-/-*^ BMSCs overexpressed with ALPL and mutant α2δ subunit. No significant change of Ca_V_1.2 or Ca_V_1.3 in cytoplasm was found in *alpl*^*-/-*^ BMSCs, *alpl*^*-/-*^ BMSCs overexpressed with ALPL and mutant α2δ subunit, *alpl*^*-/-*^ BMSCs overexpressed with ALPL and α2δ subunit. (f) α2δ immunoprecipitated ALPL. Left panel showed the expression of ALPL and right panel showed the level of ALPL following immunoprecipitation with anti-α2δ antibody in *alp/*^*f/f*^ and *alpl*^*-/-*^ BMSCs. β-actin was used as a protein loading control. Representative results were from three independent experiments.

We next investigated what region of Ca_V_1.2 and Ca_V_1.3 is important for ALPL-regulated internalization. α_2_δ subunit was indicated in trafficking α1 subunits, which influences internalization of the channels (Canti et al, 2005; Tetreault et al, 2016). Floxed *alpl* mice with *Prrx1*::*Cre* mice were crossed to generate early embryonic MSC-specific (*Prrx1-alpl*^*-/-*^) conditional *alpl* knockout mice (**Fig S5b, c**). We isolated the BMSCs from *Prrx1-alpl*^*-/-*^ mice (ALPL^-/-^) and the control *alpl*^*f/f*^ littermates (control), *alpl*^*-/-*^ BMSCs showed decreased expression of ALPL and lentivirus-overexpressing ALPL (Leti-alp) treatment elevated the membrane expression of ALPL (**Fig S5d**). To explore whether ALPL interacted with α_2_δ to regulate the internalization of Ca_V_1.2 and Ca_V_1.3, we expressed ALPL with α_2_δ or mutant α_2_δ in *alpl*^*-/-*^ BMSCs and examined the membrane expression of Ca_V_1.2 and Ca_V_1.3. *alpl*^*-/-*^ BMSCs were isolated from *Prrx1-alpl*^*-/-*^ mice and showed no expression of ALPL, Ca_V_1.2 and Ca_V_1.3 on the membrane (**Fig 4d**). The membrane expression of Ca_V_1.2 and Ca_V_1.3 was increased and ALPL was co-localized with Ca_V_1.2 and Ca_V_1.3 after transfected with ALPL and α_2_δ (**Fig 4d**). However, the membrane expression of Ca_V_1.2 and Ca_V_1.3 was not increased and ALPL was not co-localized with Ca_V_1.2 and Ca_V_1.3 after transfected with ALPL and mutant α_2_δ (**Fig 4d**). Western blot analysis also confirmed that the membrane expression of Ca_V_1.2 and Ca_V_1.3 was increased after transfected with ALPL and α_2_δ (**Fig 4e**). However, no membrane expression of Ca_V_1.2 and Ca_V_1.3 was found in *alpl*^*-/-*^ BMSCs and *alpl*^*-/-*^ BMSCs transfected with ALPL and mutant α_2_δ (**Fig 4e**). Immunoprecipitation using a control antibody did not isolate either protein in WT and *alpl*^*-/-*^ BMSCs, but an immunoprecipitation with anti-α2δ resulted in a coimmunoprecipitation with ALPL in *alpl*^*f/f*^ BMSCs (**Fig 4f**). To confirm that ALPL interacts with α_2_δ subunits and thus regulates the lineage differentiation of BMSCs, we transfected *alpl*^*+/-*^ BMSCs with α_2_δ or mutant α_2_δ and assessed their osteogenic or adipogenic induction. *alpl*^*+/-*^ BMSCs transfected with mutant α_2_δ showed decreased osteogenic differentiation and increased adipogenic differentiation compared to *alpl*^*+/-*^ BMSCs transfected with α_2_δ (**Fig S5e-h**).

### ALPL deficiency caused aberrant lineage differentiation of BMSCs through Wnt/β-catenin Pathway

In order to examine how ALPL-deficiency-induced reduction of Ca^2+^ influx affects the osteogenic and adipogenic differentiation of BMSCs, we analyzed three Ca^2+^ downstream pathways, including PKC/Erk, PI3K/Akt/GSK3β, and CaMKII/calcineurine A cascades, which are closely linked to Ca^2+^-associated regulation of osteogenic differentiation. We found that the expression level of p-Akt significantly decreased along with the reduction of p-GSK3β in *alpl*^*+/-*^ BMSCs and BMSCs transfected with shALP (**Fig 5a**). However, PKC/Erk and CaMKII/calcineurine A pathways were not changed significantly in *alpl*^*+/-*^ BMSCs and BMSCs transfected with shALP compared to WT BMSCs (**Fig S6a**). Because the decrease of GSK3β phosphorylation inhibits the nuclear translocation of β-catenin, which regulates osteogenic and adipogenic differentiation of BMSCs, we examined the expression levels of total and active β-catenin. We found that the expression of active β-catenin was decreased in both *alpl*^*+/-*^ BMSCs and BMSCs transfected with shALP (**Fig 5a**). Moreover, when we overexpressed ALPL in *alpl*^*+/-*^ BMSCs, the expression of p-Akt, p-GSK3β and active β-catenin was all increased to the level similar to WT BMSCs (**Fig 5b**). When we overexpressed Ca_V_1.2 or Ca_V_1.3 in *alpl*^*+/-*^ BMSCs, the expression of p-Akt, p-GSK3β and active β-catenin was all increased (**Fig 5c**). In addition, expression of DN-Dyn1 in *alpl*^*+/-*^ BMSCs also increased the expression of p-Akt, p-GSK3β and active β-catenin compared to *alpl*^*+/-*^ BMSCs (**Fig 5d**). Altogether, these data indicate that ALPL regulates Ca^2+^ influx to affect p-Akt and p-GSK3β expression, and subsequently target on the Wnt/β-catenin pathway in BMSCs.

**Figure 5.**
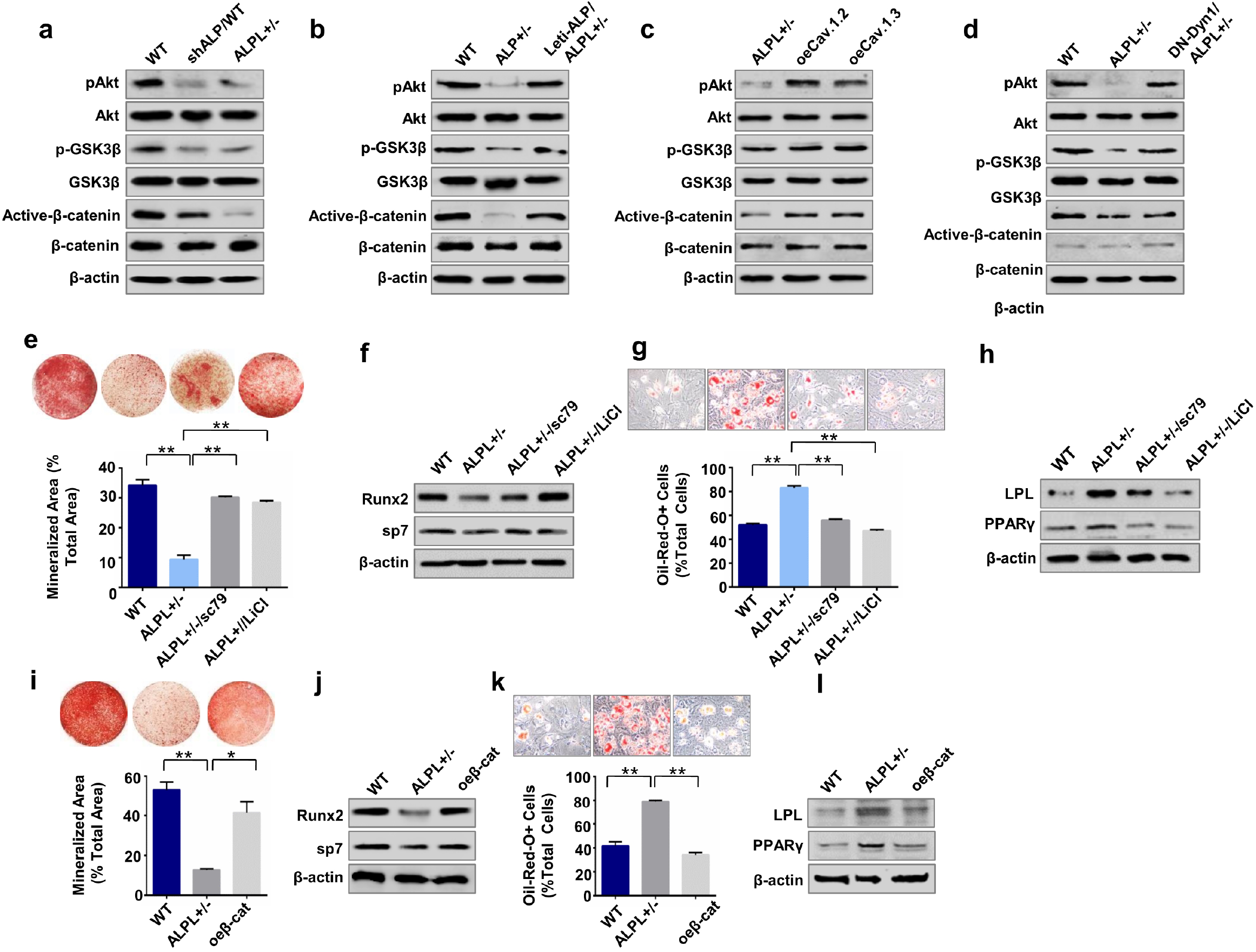
ALPL deficiency caused aberrant lineage differentiation of BMSCs through Wnt/β-catenin Pathway. (a) *alpl*^*+/-*^ and shALP BMSCs showed significant downregulation of p-AKT, p-GSK3β and active β-catenin compared to WT BMSCs. β-actin was used as a protein loading control. (b) Leti-alp/*alpl*^*+/-*^ BMSCs showed significant upregulation of p-AKT, p-GSK3β and active β-catenin compared to *alpl*^*+/-*^ BMSCs. (c) *alpl*^*+/-*^ BMSCs transfected with oeCa_V_1.2 or oeCa_V_1.3 showed significant upregulation of p-AKT, p-GSK3β and active β-catenin.(d) *alpl*^*+/-*^ BMSCs transfected with DN-Dyn1 showed significant upregulation of p-AKT, p-GSK3β and active β-catenin. (e, f) Alizarin red staining showed that *alpl*^*+/-*^ BMSCs treated by 10 μM sc79 or 10 mM LiCl had an increased capacity to form mineralized nodules when cultured under osteo-inductive conditions (e) and upregulation of the osteogenic related proteins RUNX2 and Sp7 (f). (g, h) *alpl*^*+/-*^ BMSCs treated by sc79 or LiCl showed a decreased number of Oil red O-positive adipocytes when cultured under adipo-inductive conditions (g) and downregulation of the adipogenic related proteins PPARγ2 and LPL, as assessed by western blot (h). (i, j) Alizarin red staining showed that *alpl*^*+/-*^ BMSCs transfected with plasmid-overexpressed β-catenin (oeβ-cat/ALPL+/-) had an increased capacity to form mineralized nodules when cultured under osteo-inductive conditions (i) and upregulation of the osteogenic related proteins RUNX2 and Sp7 (j). (k, l) *alpl*^*+/-*^ BMSCs transfected with plasmid-overexpressed β-catenin (oeβ-cat/ALPL+/-) showed a decreased number of Oil red O-positive adipocytes when cultured under adipo-inductive conditions (k) and downregulation of the adipogenic related proteins PPARγ2 and LPL, as assessed by western blot (l). Representative results were from three independent experiments. Error bars represent the s.d. from the mean values. *P<0.05, ** P < 0.01.

To further elucidate whether ALPL regulates the osteogenic and adipogenic differentiation through the Akt/GSK3β/Wnt/β-cantenin pathway, we used activators for Akt phosphorylation (sc79) and GSK3β phosphorylation (LiCl) to treat *alpl*^*+/-*^ BMSCs and found that sc79 and LiCl treatment increased the osteogenic differentiation, as evidenced by increased mineralized nodule formation and expression of RUNX2 and Sp7 (**Fig 5e, f**). In contrast, sc79 and LiCl treatment decreased adipogenic differentiation of *alpl*^*+/-*^ BMSCs, as assessed by Oil red O staining to show decreased number of adipocytes and western blot to show downregulation of PPARγ2 and LPL under adipogenic culture conditions (**Fig 5g, h**). Moreover, we also overexpressed β-catenin (oeβ-cat) in *alpl*^*+/-*^ BMSCs (**Fig S6b**) and found rescued lineage differentiation in *alpl*^*+/-*^ BMSCs, as evidenced by the increased osteogenic differentiation and decreased adipogenic differentiation (**Fig 5i-l**).

### Raising the intracellular level of calcium by ionomycin rescued ALPL deficiency-induced age-related osteoporosis

Ionomycin was reported to cause a robust rise of Ca^2+^ influx and may inhibited calcium channel endocytosis (Green et al, 2007). We confirmed that ionomycin treatment increased the calcium influx and the membrane expression of Ca_V_1.2 and Ca_V_1.3 of *alpl*^*+/-*^ BMSCs (**Fig S7a, b**). MicroCT and histological analyses showed that bone mineral density (BMD), bone volume/total volume (BV/TV), and distal femoral trabecular bone structure (Tb.N) in 3-month-old *alpl*^*+/-*^ mice were markedly decreased compared to the control WT littermates (**Fig. 6a, b**). We treated *alpl*^*+/-*^ mice with ionomycin, which caused a robust rise of Ca^2+^ influx in cells (Green et al, 2007). MicroCT and histological analyses showed that BMD, BV/TV and Tb.N in 3-month-old *alpl*^*+/-*^ mice treated by ionomycin were markedly increased compared to the *alpl*^*+/-*^mice (**Fig. 6a, b**). To observe alteration of osteogenic/adipogenic lineage differentiation in vivo, we examined the number of adipocytes in bone marrow of WT, *alpl*^*+/-*^ mice and *alpl*^*+/-*^ mice treated by ionomycin. Interestingly, Oil red O staining showed that the number of adipocytes in *alpl*^*+/-*^ mice bone marrow was markedly increased compared to WT littermates (**Fig. 6c**), indicating that *alpl* deficiency increased adipogenic lineage differentiation. However, the number of adipocytes in *alpl*^*+/-*^ mice bone marrow after ionomycin treatment was markedly decreased compared to *alpl*^*+/-*^ mice (**Fig. 6c**). To confirm that *alpl* deficiency directly contributes to the decreased osteogenesis, a calcein double labeling analysis was used to show decreased bone formation rate in *alpl*^*+/-*^ mice but elevated bone formation rate in *alpl*^*+/-*^ mice treated by ionomycin (**Fig. 6d**).

**Figure 6.**
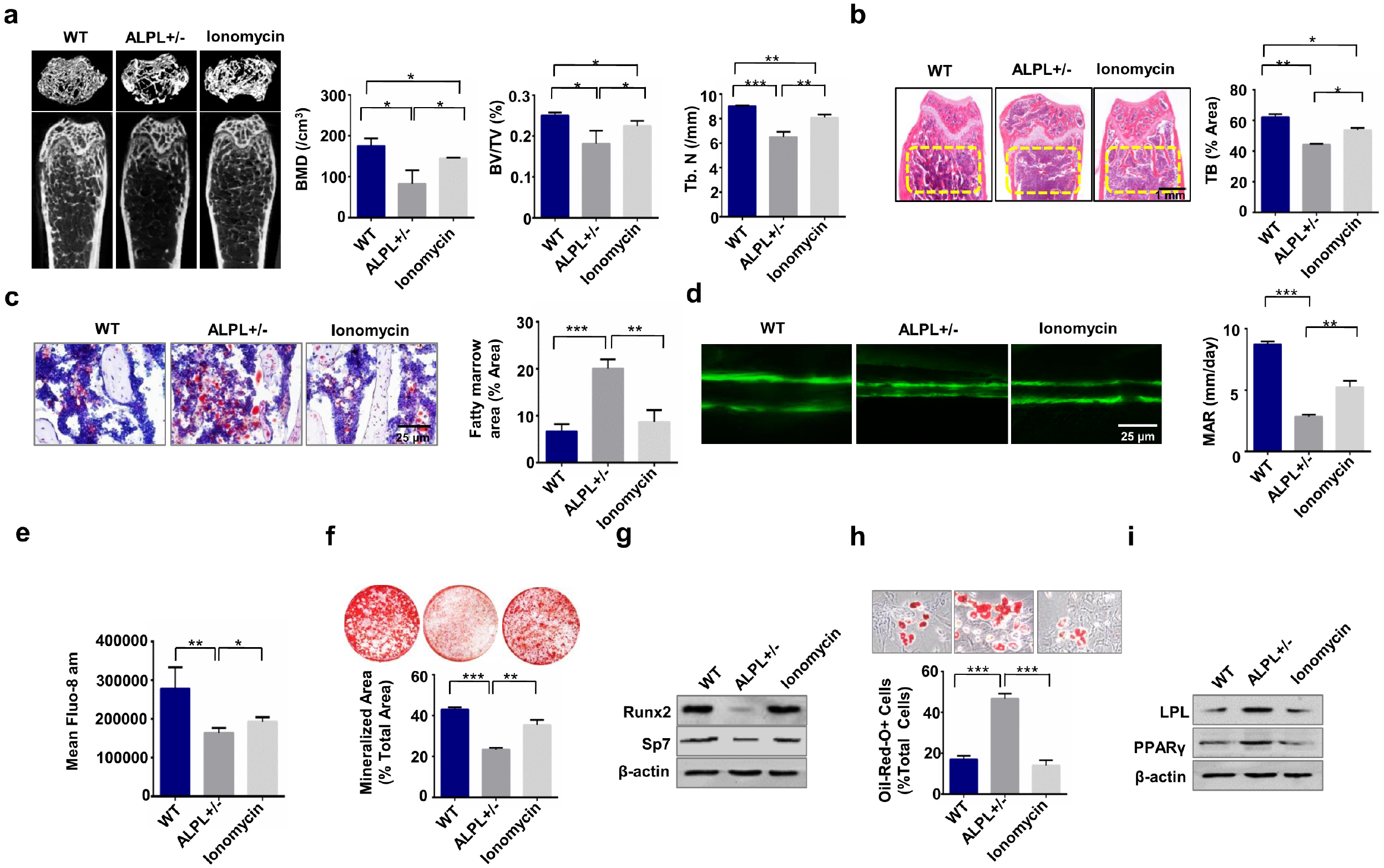
Raising the intracellular level of calcium by ionomycin rescued ALPL deficiency-induced age-related osteoporosis. (a) MicroCT analysis showed that *alpl*^*+/-*^ mice (n=12) had significantly decreased bone mineral density (BMD), bone volume over total volume (BV/TV), and trabecular number (Tb. N) in trabecular bone (TB) of the distal femur compared to *WT* littermates and *alpl*^*+/-*^ mice treated by ionomycin (n=10). (b) H&E staining showed a decreased TB volume (yellow circled area) in the distal femurs of *alpl*^*+/-*^ mice compared to the WT group and *alpl*^*+/-*^ mice treated by ionomycin. Scale bar, 1 mm. (c) Representative histological images of distal femurs showed a significantly increased number of adipocytes in *alpl*^*+/-*^ mouse bone marrow, as assessed by Oil red O staining. Scale bar, 25 μm. (d) Calcein double labeling assay showed a significantly decreased bone formation rate in *alpl*^*+/-*^ mice compared to the WT and *alpl*^*+/-*^ mice treated by ionomycin. Scale bar, 25 μm. (e) Intracellular level of Ca^2+^ in *alpl*^*+/-*^ BMSCs was decreased compared to WT BMSCs and ionomycin treatment elevated the intracellular level of Ca^2+^ in *alpl*^*+/-*^ BMSCs. (f) Alizarin red staining showed that *alpl*^*+/-*^ BMSCs had a decreased capacity to form mineralized nodules compared to WT BMSCs when cultured under osteo-inductive conditions. Ionomycin treatment of *alpl*^*+/-*^ BMSCs showed an elevated capacity to form mineralized nodules. (g) Western blot analysis showed that *alpl*^*+/-*^ BMSCs had decreased expression of osteogenic related proteins RUNX2 and Sp7 compared to WT BMSCs and ionomycin treatment of *alpl*^*+/-*^ BMSCs. β-actin was used as a protein loading control. (h, i) *alpl*^*+/-*^ BMSCs showed an increased number of Oil red O-positive cells when cultured under adipo-inductive conditions (h) and upregulation of the adipogenic related proteins PPARγ2 and LPL, as assessed by western blot (i) compared to WT BMSCs and ionomycin treatment of *alpl*^*+/-*^ BMSCs. Representative results were from three independent experiments. Error bars represent the s.d. from the mean values. *P<0.05, ** P < 0.01. *** P < 0.001.

Moreover, we found that intracellular level of Ca^2+^ in *alpl*^*+/-*^ BMSCs was decreased compared to WT BMSCs and ionomycin treatment elevated the intracellular level of Ca^2+^ in *alpl*^*+/-*^ BMSCs (**Fig. 6e**). We found impaired osteogenic differentiation and increased adipogenic differentiation of *alpl*^*+/-*^ BMSCs, as evidenced by decreased mineralized nodule formation and elevated numbers of adipocytes (**Fig. 6f, h**). The deceased expression of the osteogenic markers RUNX2 and Sp7 and increased expression of the adipogenic regulators PPARγ2 and LPL were assayed by western bolt (**Fig. 6g, i**). Ionomycin treatment increased the osteogenic differentiation and decreased adipogenic differentiation of *alpl*^*+/-*^ BMSCs (**Fig. 6f-i**).

### Ionomycin treatment rescued osteoporosis in BMSC-specific conditional *alpl* knockout mice

To further confirm whether ALPL deficiency in BMSCs caused altered osteogenesis and adipogenesis *in vivo*, we found BV/TV, and Tb.N in 3-month-old *Prrx1-alpl*^*-/-*^ mice were markedly decreased compared to the control *alpl*^*f/f*^ littermates (**Fig. 7a, b**). Floxed *alpl* littermates (*alpl*^*f/f*^) were used as the controls. MicroCT and histological analyses showed that BMD, Moreover, BMD, BV/TV, and Tb.N in 3-month-old *Prrx1-alpl*^*-/-*^ mice treated by ionomycin were markedly increased compared to the *Prrx1-alpl*^*-/-*^ mice (**Fig. 7a, b**). To further observe alteration of osteogenic/adipogenic lineage differentiation in BMSCs, we examined the number of adipocytes in bone marrow of *alpl*^*f/f*^, *Prrx1-alpl*^*-/-*^ mice and *Prrx1-alpl*^*-/-*^ mice treated by ionomycin and found that Oil red O staining showed that the number of adipocytes in *Prrx1-alpl*^*-/-*^ bone marrow was markedly increased compared to control *alpl*^*f/f*^ littermates (**Fig. 7c**). However, the number of adipocytes in *Prrx1-alpl*^*-/-*^ bone marrow after ionomycin treatment was markedly decreased compared to *Prrx1-alpl*^*-/-*^ mice (**Fig. 7c**). Calcein double labeling analysis showed decreased bone formation rate in *Prrx1-alpl*^*-/-*^ mice relative to control *alpl*^*f/f*^ mice (**Fig. 7d**). Ionomycin treatment reversed the impaired osteogenesis in *Prrx1-alpl*^*-/-*^ mice. Moreover, the serum levels of RANKL and OPG were not significantly changed, as assessed by ELISA (**Fig S7c, d**), suggesting that osteoclasts may be not altered in *Prrx1-alpl*^*-/-*^ mice. Intracellular level of Ca^2+^ in *alpl*^*-/-*^ BMSCs was decreased compared to control BMSCs and ionomycin treatment elevated the intracellular level of Ca^2+^ in *alpl*^*-/-*^ BMSCs (**Fig. 7e**). In addition, BMSCs from *Prrx1-alpl*^*-/-*^ mice showed decreased osteogenic differentiation and increased adipogenic differentiation compared to BMSCs from *alpl*^*f/f*^ mice (**Fig. 7f-i**). BMSCs from *Prrx1-alpl*^*-/-*^ mice treated by ionomycin showed increased osteogenic and dereased adipogenic differentiation (**Fig. 7f-i**). These results indicated that ALPL deficiency in BMSCs induces age-related osteoporosis phenotype and ionomycin treatment reversed the osteoporosis phenotype.

**Figure 7.**
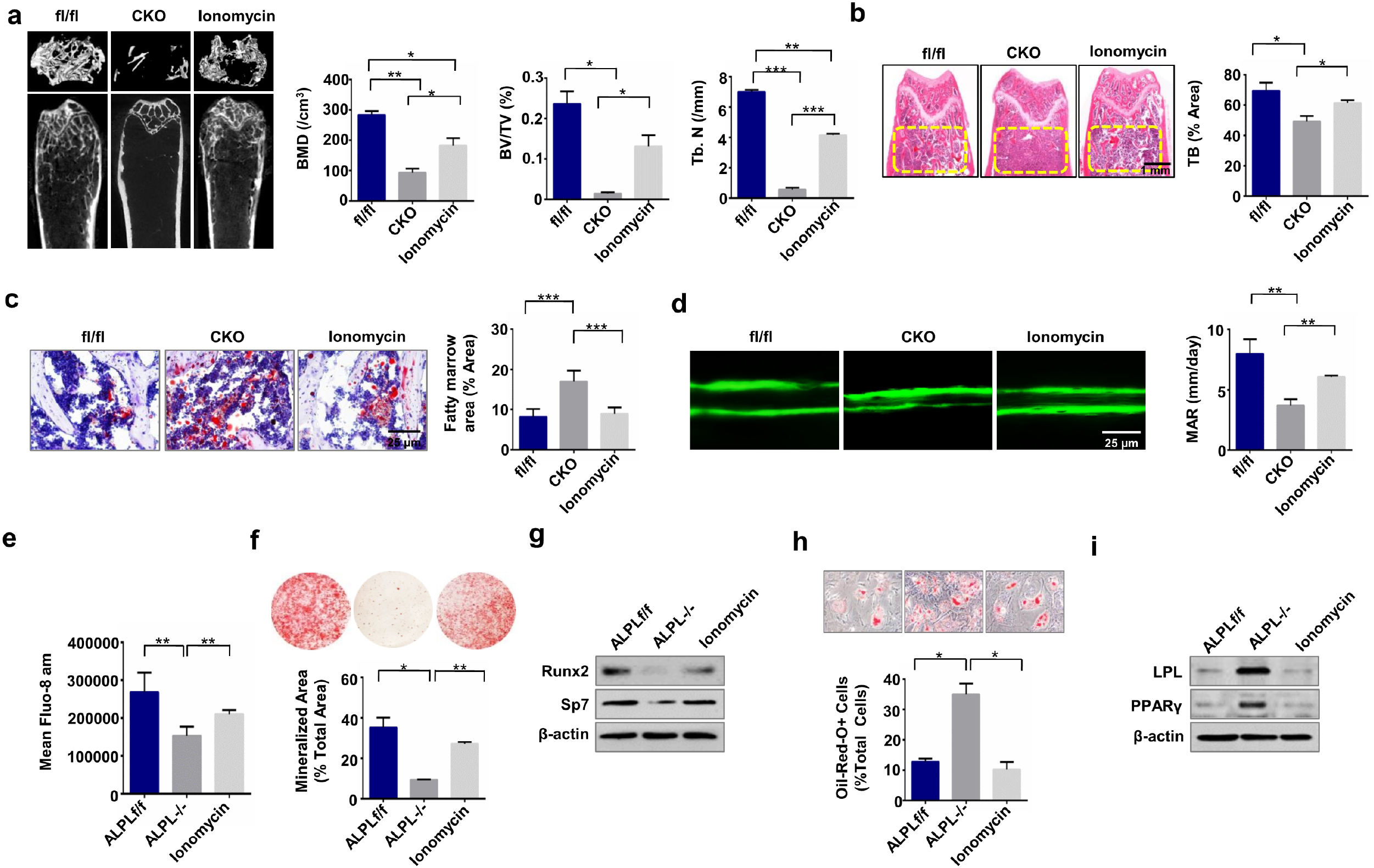
Ionomycin treatment rescued osteoporosis in BMSC-specific conditional *alpl* knockout mice. (a) MicroCT analysis showed that *Prrx1-alpl*^*-/-*^ mice (n=12) had significantly decreased bone mineral density (BMD), bone volume over total volume (BV/TV), and trabecular number (Tb. N) in trabecular bone (TB) of the distal femur compared to *alpl*^*f/f*^ littermates and *Prrx1-alpl*^*-/-*^ mice treated by ionomycin (n=10). (b) H&E staining showed a decreased TB volume (yellow circled area) in the distal femurs of *Prrx1-alpl*^*-/-*^ mice compared to the *alpl*^*f/f*^ control group and *Prrx1-alpl*^*-/-*^ mice treated by ionomycin. Scale bar, 1 mm. (c) Representative histological images of distal femurs showed a significantly increased number of adipocytes in *Prrx1-alpl*^*-/-*^ mouse bone marrow, as assessed by Oil red O staining. Scale bar, 25 μm. (d) Calcein double labeling assay showed a significantly decreased bone formation rate in *Prrx1-alpl*^*-/-*^ mice compared to the *alpl*^*f/f*^ controls and *Prrx1-alpl*^*-/-*^ mice treated by ionomycin. Scale bar, 25 μm. (e) Intracellular level of Ca^2+^ in *alpl*^*-/-*^ BMSCs was decreased compared to WT BMSCs and ionomycin treatment elevated the intracellular level of Ca^2+^ in *alpl*^*-/-*^ BMSCs. (f) Alizarin red staining showed that *alpl*^*-/-*^ BMSCs had a decreased capacity to form mineralized nodules compared to WT BMSCs when cultured under osteo-inductive conditions. BMSCs from *Prrx1-alpl*^*-/-*^ mice treated by ionomycin showed an elevated capacity to form mineralized nodules. (g) Western blot analysis showed that *alp*^*-/-*^ BMSCs had decreased expression of osteogenic related proteins RUNX2 and Sp7 compared to WT BMSCs and ionomycin treatment of *alpl*^*-/-*^ BMSCs. β-actin was used as a protein loading control. (h, i) *alpl*^*-/-*^ BMSCs showed an increased number of Oil red O-positive cells when cultured under adipo-inductive conditions (h) and upregulation of the adipogenic related proteins PPARγ2 and LPL, as assessed by western blot (i) compared to WT BMSCs and ionomycin treatment of *alpl*^*-/-*^ BMSCs. Representative results were from three independent experiments. Error bars represent the s.d. from the mean values. *P<0.05, ** P < 0.01. *** P < 0.001.

### ALPL deficiency promoted the internalization of L-type Ca^2+^ channel in HPP patient derived BMSCs

We also collected bone marrow BMSCs from two HPP patients with mutation in ALPL gene (A1 and A2, **Table S1**). The expression of ALPL on membrane and in cytoplasm was decreased in BMSCs from two HPP patients compared to normal human bone marrow BMSCs (control) (**Fig 8a**). We further confirmed that whether lack of ALPL results in decreased KCl-induced Ca^2+^ influx in A1 and A2 BMSCs. KCl-induced Ca^2+^ influx was significantly decreased in culture-expanded A1 and A2 BMSCs compared to control BMSCs (**Fig 8b**) and overexpression of ALPL elevated KCl-induced Ca^2+^ influx (**Fig 8c, d**) in A1 and A2 BMSCs. Moreover, we also found that expression of DN-Dyn1 prevented the decrease of KCl-induced Ca^2+^ influx in A1 and A2 BMSCs (**Fig 8c, d**). Overexpression of ALPL or expression of DN-Dyn1 in A1 and A2 BMSCs increased the membrane expression of Ca_V_1.2 and Ca_V_1.3 as the confocal images showed (**Fig 8e**), suggesting that ALPL deficiency causes channel internalization of Ca_V_1.2 and Ca_V_1.3 in human model. To confirm that ALPL regulates the lineage differentiation of BMSCs, we overexpressed ALPL in A1 and A2 BMSCs and assessed their osteogenic or adipogenic ablity after induction. A1 and A2 BMSCs transfected with ALPL vector showed increased osteogenic differentiation and decreased adipogenic differentiation (**Fig S8a-d**). All of the above data show that ALPL maintains lineage differentiation of MSCs through directly binding to α2δ subunit of L-type Ca^2+^ channels and inhibiting the internalization of L-type Ca^2+^ channels, thus increases Ca^2+^ influx (**Fig S9**).

**Figure 8.**
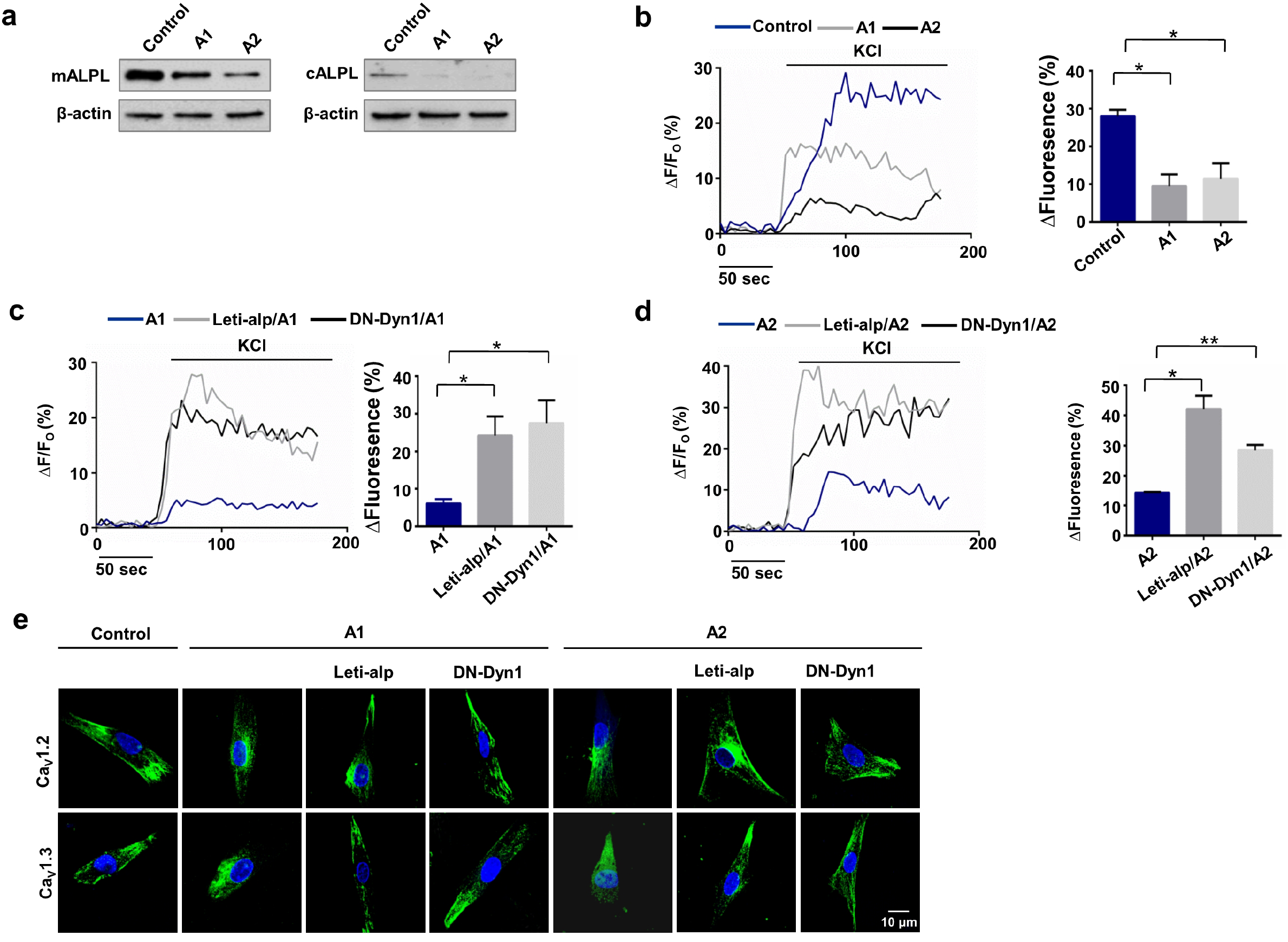
ALPL deficiency promoted the internalization of L-type Ca^2+^ channel in HPP patient derived BMSCs. (a) The expression of ALPL on membrane and in cytoplasm was decreased in BMSCs from two HPP patients compared to normal human BMSCs. β-actin was used as a protein loading control. (b) Intracellular Ca^2+^ imaging analysis showed that KCl-induced Ca^2+^ influx was significantly decreased in cultured BMSCs from HPP patients compared to normal human BMSCs. (c) Overexpression of ALPL or transfected with DN-Dyn1 in BMSCs from A1 patient showed elevated KCl-induced Ca^2+^ influx. (d) Overexpression of ALPL or transfected with DN-Dyn1 in BMSCs from A2 patient showed elevated KCl-induced Ca^2+^ influx. (e) Representative images of confocal laser scanning microscope showed membrane location of Ca_V_1.2 and Ca_V_1.3 in control BMSCs, A1 BMSCs, A2 BMSCs, A1 and A2 BMSCs overexpressed with ALPL, A1 and A2 BMSCs transfected with DN-Dyn1. Scale bar, 10 μm. Representative results were from three independent experiments. Error bars represent the s.d. from the mean values. *P < 0.05. **P < 0.01.

## Discussion

The deficiency of HPP could be ameliorated by bone anabolic and/or enzyme replacement treatment strategies thus far. Bone anabolic treatment such as recombinant human parathyroid hormone (PTH) analogs showed controversial treatment efficacy (Seefried et al, 2017). Asfotase alfa (Strensiq, Alexion), a bone-targeted enzyme replacement therapy, was approved for long-term treatment of pediatric-onset HPP in the United States, Europe, Canada, and Japan. However, there are no guidelines for selecting adult patients for treatment, for evaluating the results of treatment, or determining the optimal duration of treatment at this time. These patients also can develop secondary osteoporosis, bone marrow edema, and delayed fracture healing or difficulties with implant failure (Seefried et al, 2017). Thus, there is an urgent need for identifying further bone targeted treatment options for such adult HPP patients.

Our previous study showed that Alpl deficiency results in premature bone ageing characterized by bone mass loss and parallel marrow fat gain. Although a pivotal role for ALPL in skeletal matrix mineralization is established, the mechanism of differentiation of BMSCs regulated by ALPL remains uncertain. Hypercalcemia was found in severely affected infantile HPP, which suggests that ALPL may modulate calcium homeostasis. In this study, we found that raising the intracellular level of calcium in BMSCs by ionomycin rescued ALPL deficiency caused age-related osteoporosis, which suggest that targeting the calcium channel is a new approach for adult HPP treatment. Moreover, our study showed a new function of ALPL in controlling the Ca^2+^ influx through regulating the internalization of calcium channels, which balanced the osteogenic and adipogenic differenation of BMSCs.

Voltage-dependent calcium channel are an important route for the entry of Ca^2+^ into cells upon membrane depolarization to regulate the calcium homeostasis. Previous study showed that VGCCs in BMSCs and osteoblast regulate bone formation and manipulating VGCCs promotes bone repair (Sun et al, 2012; Zhang et al, 2016). Of note, the long-lasting voltage-gated calcium channel (L-VGCC), a major channel of calcium influx, is a part of the high-voltage activated family of voltage-gated calcium channel. L-type calcium channel is considered to play an important role in regulating BMSCs function (Barradas et al, 2012; Wen et al, 2012). L-VGCCs, which consist of four subunits (Ca_V_1.1, Ca_V_1.2, Ca_V_1.3 and Ca_V_1.4), are expressed and distributed in different tissues (Snutch & Reiner, 1992). Ca_V_1.2 and Ca_V_1.3 constitute the major fraction of L-type calcium channels in mammals (Hell et al, 1993; Ludwig et al, 1997; Takimoto et al, 1997). However, the regulation of Ca_V_1.2 and Ca_V_1.3 in BMSCs are still unknown. Here we found that lack of ALPL caused decreased number of Ca_V_1.2 and Ca_V_1.3 in the membrane and decreased Ca^2+^ influx in BMSCs, which leads to the aberrant lineage differentiation of BMSCs.

The involvement of ALPL in channel internalization leads to a change in the number of Ca_V_1.2 and Ca_V_1.3 at the cell surface. Although several other proteins have been found to regulate the membrane expression of VGCCs, including calmodulin, Akap15, Akap9 and eIF3e, and these associations play an important role in connecting VGCCs and intracellular signaling pathways (Green et al, 2007). Of these proteins, ALPL is unusual in that it is an ectoenzyme, which hydrolyzes several substrates. Our study showed a new function of ALPL in controlling the Ca^2+^ influx through regulating the internalization of Ca_V_1.2 and Ca_V_1.3 in BMSCs. The results suggest that ectoenzymes on the cell membrane may bind channels in the cell and regulate the trafficking of channels. Precisely how ALPL regulates L-type Ca^2+^ channel trafficking is unclear. Our co-localization data indicates that there is a direct contact between ALPL and L-type Ca^2+^ channel. α2δ subunit of L-type Ca^2+^ channel is responsible for regulating the trafficking of channels (Canti et al, 2005; Tetreault et al, 2016). We further demonstrate that ALPL may bind to the α2δ subunit to regulate L-type Ca^2+^ channel trafficking, as the mutation of α2δ subunit prevents the membrane expression of Ca_V_1.2 and Ca_V_1.3. The conditional binding of ALPL to Ca_V_1.2 and Ca_V_1.3 suggests that the composition of Ca_V_1.2 and Ca_V_1.3 protein complex with ALPL may be an important factor in channel regulation.

Skeletal defects in HPP include rickets, osteomalacia, fractures, bone pain and various dental defects (Foster et al, 2013; Foster et al, 2014). To understand the physiological role of ALPL and evaluate the potential treatments, several lines of ALPL knock-out mouse were generated (Narisawa et al, 1997; Waymire et al, 1995). Homozygous mice show severe bone disease, but they often die before puberty (Foster et al, 2017). However, here we found that ALPL deficiency in BMSCs caused decreased osteogenic differentiation and increased adipogenic differentiation. *alpl*^*+/-*^ mouse model phenocopies the adult HPP, and these mice showed inhibited bone formation but increased adipose tissue in the bone matrix. Moreover, the bone formation was inhibited when we generate *alpl* conditional knockout mice of BMSCs, which was consistent with the recent reports (Foster et al, 2017). Recent data have shown that increased marrow adipose tissue is correlated with dysfunction of bone and hematopoietic regeneration (Ambrosi et al, 2017). We also found that bone formation was inhibited but the adipose tissue was increased in the bone matrix of *alpl* conditional knockout mice, which suggests that ALPL regulates the osteogenic/adipogenic differentiation of BMSCs and causes the osteoporosis of HPP.

Our finding of the involvement of ALPL in calcium homeostasis revealed the molecular mechanism underlining the balance of BMSCs osteogenic and adipogeninc differentiation. The change of Ca^2+^ influx in BMSCs by H_2_S regulates the osteogenic differentiation through PCK/Erk/Wnt pathway (Liu et al, 2014). Here we demonstrated that Akt/GSK3β/Wnt/β-cantenin pathway was the downstream of ALPL regulating Ca^2+^ influx. Akt/GSK3β/Wnt/β-cantenin pathway further balanced the osteogenic and adipogenic differentiation of BMSCs and the bone formation. In our study, we found that ALPL was required to maintain intracellular Ca^2+^ influx by regulating internalization of L-type Ca^2+^ channel *via* binding to the α2δ subunits in BMSCs. Altered intracellular Ca^2+^ influx caused by ALPL deficiency results in osteoporotic phenotype due to downregulated osteogenic differentiation and upregulated adipogenic differentiation in both human and mouse BMSCs. Inhibition of calcium channel internalization by ionomycin increased the calcium influx enhanced the bone formation of *alpl*^*+/-*^ mice and *Prrx1-alpl*^*-/-*^ mice, suggesting that targeting calcium channel internalization is a potential treatment strategy for adult HPP patients.

## Materials and Methods

### Mice

C57BL/6J and B6.129S7-*Alpl*^*tm1Sor*^/J mice were purchased from Jackson Laboratory (Bar Harbor, ME, USA). We produced mice that were homozygous for ALPL^flox^ using CRISPR/Cas9 system, an ALPL allele in which the DNA segment includes exons 3 and 4 was flanked by *loxP* sites. By coinjection of Cas9/sgRNAs and ALPL targeting vector into zygote, we achieved ALPL-floxed heterozygous mice. To generate a tissue-specific Cre-mediated ALPL knockout model, Prrx1-Cre (C57BL/6-Prrx1^*tm1(iCre*)^/Bcgen (Beijing Biocytogen Co., Ltd. Beijing, China) mice were mated with mice heterozygous for the ALPL-floxed allele. The offspring inherited Prrx1-Cre and two ALPL-floxed alleles. Routine mouse genotyping was performed by PCR. The following primer pairs were used for Cre and floxed alleles: Prrx1^iCre^ primer 1: AGCGTTTGGTGTTGATTCGAGC; Prrx1^iCre^ primer 2: AGTCCCGGTGACTCCAGCAG; Prrx1^iCre^ primer 3: TGGTGCACAGTCAGCAGGTTG; ALPL-3′loxP-F: CCTGCTACCTGCTTGCTGGCAATGG; ALPL-3′loxP-R: AGGACACCAAAGACCAGGGACACTA. All animal experiments were performed under institutionally approved protocols for the use of animal research (University of Fourth Military Medical University protocol number k9-024).

### Isolation of mouse bone marrow mesenchymal stem cells

Bone marrow cells were flushed from the bone cavities of mouse femurs and tibias with 2% heat-inactivated fetal bovine serum (FBS; Hangzhou Sijiqing Biological Engineering Materials Co., Ltd. Zhejiang, China) in PBS. A single-cell suspension of all nucleated cells was obtained by passing all bone marrow cells through a 70 μm cell strainer (BD Bioscience, NJ, USA). All the single cells (1 × 10^6^) were seeded in 100 mm culture dishes (Corning, NY, USA) and initially incubated for 48 h at 37 °C and 5% CO_2_. To eliminate the non-adherent cells, the cultures were washed with PBS twice on the second day. The attached cells were cultured for 16 days with alpha minimum essential medium (α-MEM, Gibco BRL, Gaithersburg, MD, USA) supplemented with 20% FBS, 2 mM l-glutamine (Invitrogen Life Technology, Carlsbad, CA, USA), 55 μM 2-mercaptoethanol (Invitrogen), 100 U ml^-1^ penicillin, and 100 μg ml^-1^ streptomycin (Invitrogen). To confirm mesenchymal stem cell character, we used flow cytometric analysis to show that these BMSCs were positive for Sca-1, CD44 and CD73, and negative for CD34 and CD45 (BD Biosciences, San Diego, CA, USA).

### Calcium imaging

Calcium imaging was performed using the confocal laser microscopy (Zeiss, Oberkochen FV1000, Germany). Intracellular Ca^2+^ level ([Ca^2+^]_i_) was represented by the Fluo-3 fluorescence intensity as described previously (Merritt et al, 1990). Briefly, BMSCs were cultured in 12-well plate and incubated with 5 μM Fluo-3/AM dye (Invitrogen, Life Technology, Carlsbad, CA, USA) for 30 min in α-MEM at 37 °C. BMSCs were again washed for three times with calibrated EGTA/Ca^2+^ solutions. KCl (30 mM) or TG (20 μM, Sigma-Aldrich, St. Louis., MO, USA) was added to test which type of calcium channel was affected. Images were collected every 4 s at 2 Hz with excitation at 488 nm and emission at 530 nm. Data are presented as Fluo-3 fluorescence intensity increase ratio: R =*ΔF/F_0_*, where *ΔF*= *F–F_0_.* F is the fluorescence value detected and F_0_ is the minimum fluorescence value.

### Confocal Microscopy

Confocal images were acquired with a Zeiss Oberkochen FV1000 confocal laser scanning microscope, using a×60 oilimmersion objective. BMSCs were fixed with 3.7% paraformaldehyde in distilled water at 4°C for 10 min, incubated overnight with primary antibodies, followed by secondary antibodies for 1 hr. The nucleuses were stained with 1 µg/ml Hoechst 33342. Images were acquired using an argon laser (excitation, 488 nm; emission, BP505–530 nm emission filter) for FITC-labeled-Cav1.2 or Cav1.3, a UV laser for excitation and a BP385–470 nm emission filter for Hoechst 33342, and a He-Ne laser (excitation, 543 nm; emission filter, LP650nm) for Cy3-labeled-ALPL. 1× 10^5^ BMSCs were plated onto coverslips, the cells were treated with 10 μM ATP or 10 μM ppi or 1μg/mL ionomycin for 1 h at the next day, and then for immunofluorescence staining of Cav1.2 or Cav1.3. Plasma membrane localization of Ca_V_1.2 or Ca_V_1.3 in BMSCs stained with anti-Ca_V_1.2 or anti-Ca_V_1.3 antibody was recorded for more than 10 cells. Co-localization of L-type Ca^2+^ channel (Ca_V_1.2 or Ca_V_1.3) and ALPL were also observed using laser scanning confocal microscope with images obtained using FV10-ASW Viewer 4.2 (Zeiss, OberkochenFV1000, Germany).

To record the time-course change of internalization of Ca_V_1.2, plasmid encoding DsRed-Cav1.2 was generated and transfected into the BMSCs. The plasmid was constructed by introducing a DsRed segment into the Ca_V_1.2 plasmid (Addgene plasmid # 26572) according to the instruction of the ClonExpress® II One Step Cloning Kit (Vazyme, Nanjing, China). Dio (Thermo Fisher Scientific, MA, USA) was used to label the cell membrane and DAPI (Thermo Fisher Scientific) was used to label nucleus. Average co-localization intensity was determined by selecting a region of interest (ROI) corresponding to the cell’s footprint in the first image and measuring the average intensity in that region over the entire time course. ROI was visually selected in a region of the Dio labeled plasma membrane. Cells that moved out of this ROI were excluded from analysis. The dwell time of channels at the membrane was measured for more than 15 ROIs from at least five cells per condition. We recorded the time dependent red fluorescence intensity of these regions. Quantitative data was presented as fluorescence intensity increase ratio: *R =F/F_0_*. *F* is the fluorescence value detected and *F*_*0*_ is the minimum fluorescence value.

### Cell Surface Protein Isolation

The expression level of Ca^2+^ channels in the surface membrane was determined by isolating cell surface proteins using Membrane and Cytosol Protein Extraction Kit (Beyotime, Shanghai, China). The cells were lysed using lysis buffer containing 1 mM EDTA, 1 mM dithiothreitol, complete protease inhibitors and 1 mM PMSF. The homogenate was centrifuged for 10 min at 700g to remove the nuclear fraction. The supernatant was then centrifuged for 30 min at 14,000g and 4°C to obtain the cytoplasmic and membrane fractions. The protein concentrations were measured by the Bradford Protein Assay Kit (Beyotime, Shanghai, China). Then equal amounts of the cytoplasmic and membrane proteins were saved as direct input for immunoblot experiments.

### Osteogenic differentiation

BMSCs were cultured under osteogenic culture conditions containing 2 mM β-glycerophosphate (Sigma-Aldrich, St. Louis., MO, USA), 100 μM L-ascorbic acid 2-phosphate (MP Biomedicals, Irvine, CA, USA), and 10 nM dexamethasone (Sigma-Aldrich, St. Louis., MO, USA) in the growth medium. Two weeks after induction, staining was performed to detect matrix mineralization with 1% Alizarin Red S (Sigma-Aldrich) for 3 min at room temperature. 10 μM SC79 or 10 mM LicL were added to the osteogenic culture medium for induction. The stained areas were quantified using NIH Image J software and shown as a percentage of the total area.

### Adipogenic differentiation

For adipogenic induction, 500 nM isobutylm ethylxanthine (MP Biomedicals, Irvine, CA, USA), 60 μM indomethacin (Sigma-Aldrich), 500 nM hydrocortisone (MP Biomedicals, Irvine, CA, USA), 10 μg/ml insulin (Sigma-Aldrich), and 100 nM L-ascorbicacid phosphate were added into the growth medium. 10 μM SC79 or 10 mM LicL were added to the adipogenic culture medium for induction. After 7 days, the cultured cells were stained with Oil Red-O (Sigma-Aldrich), and positive cells were quantified under microscopy and shown as a percentage of the total cells.

### Western immunoblotting

Cells were lysed in M-PER mammalian protein extraction reagent (Thermo, MA, USA) with protease and phosphatase inhibitors (Roche, Basel, Switzerland), and proteins were quantified using protein assays (Bio-Rad Laboratories, Shanghai, China). 20 μg of proteins were separated by SDS-PAGE and transferred to a nitrocellulose membrane (Millipore, Billerica, MA, USA). Membranes were blocked with 0.1% Tween 20 and 5% BSA for 1 h before overnight incubation with the primary antibody diluted in blocking solution. Antibody to mouse ALP was purchased from R&D Systerms, antibodies to Cav1.1, Cav1.2, Cav1.3, Cav2.1, Cav2.2, Cav3.1, Cav3.2, Cav3.3 and Ca_V_α2δ1 were purchased from alomone labs (Alomone, Jerusalem, Israel). Antibodies to mouse phospho-Erk1/2 (Thr202/Tyr204), Erk1/2, GSK-3β, phospho-GSK-3β, phospho-AKT, β-catenin, RUNX2, SP7, PPARγ, LPL, phospho-PKC, PKC, phospho-CamkII, AKT were purchased from Abcam (Cambridge, UK). Antibodies to mouse CamkII and active β-catenin were obtained from Millipore (Billerica, MA, USA). Antibody to mouse β-actin was purchased from Boster (Wuhan, China). The membranes were incubated for 1 h in HRP-conjugated secondary antibody diluted at 1:40,000 in blocking solution. Immunoreactive proteins were detected using an enhanced chemiluminescence kit (Amersham Biosciences, Piscataway, NJ, USA). The intensity of bands was measured using NIH ImageJ software and normalized to β-actin.

### Co-Immunoprecipitation

In order to test whether ALPL and L-VGCC (Cav1.2 or Cav1.3) bind each other in cells, co-immunoprecipitation was performed as previously described (J et al, 2008). The cells were completely lysed by Cell lysis buffer for Western and IP (Beyotime, Shanghai, China). The lysates were incubated with primary antibodies for overnight at 4°C and then protein A/G magnetic beads (Millipore, USA) were added for 2 h at 4°C. Immuno-complexes were washed three times with PBS containing 0.1% Tween-20 and subsequently subjected to western blot analysis.

### Plasmids construction

For shRNA knockdown experiments, we constructed shALPL lentiviral vector according to the protocol of pLKO.1-TRC Cloning Vector (Moffat et al, 2006). In brief, to generate oligos for cloning into pLKO.1, we synthesized the following oligonucleotides: forward oligo: CCGGGCAGTATGAATTGAATCGGAACTCGAGTTCCGATTCAATTCATACTGCTTTTTG, reverse oligo: AATTCAAAAAGCAGTATGAATTGAATCGGAACTCGAGTTCCGATTCAATTCATACTGC.

Then the forward and reverse oligos were annealed and ligated into the pLKO.1 vector, producing a final plasmid that expresses the shRNA of ALPL. To overexpress ALPL in BMSCs, we constructed lentiviral vector with ALPL cDNA. The ALPL (GenBank Accession No. NM_000478.5) cDNA was amplified by PCR using the primer pairs: Forward: 5’-ACTGGATCCTCCAGGGATAAAGCAGGTCT-3’; Reverse: 5’-TATCTCGAGTGGGAAGTTGGCATCTGTC-3’. And then the ALPL gene was inserted into the vector pENTR^™^2B (Invitrogen Life Technology, Carlsbad, CA, USA), a Gateway LR recombination reaction between the ALPL clone vector and pLenti6.3 / V5-DEST, to generate an expression clone Lenti-ALPL. The cDNA encoding β-catenin (GenBank Accession No. NM_007614.3) was amplified by TaKaRa LA Taq^R^ with GC Buffer (TaKaRa, Japan). Primers were as following: forward: 5′-CGGGGAGGCGGAGACGGAGCAC-3′, reverse: 5′-CCAGCCCACCCCTCGAGCCCTCTC-3′. And then the restriction enzymes SalI and BamHI were used to clone the above fragment into the backbone vector pIRES2-EGFP (Clonetech, USA).

pDsRed-Cav1.2 was constructed according to the manufacturer’s protocol of the ClonExpress Ultra One Step Cloning Kit (Vazyme, Nanjing, China). In brief, Cav1.2 was amplified by prierms: forward: 5’-GGGGTACCATGGTCAATGAAAACACGAGG-3 ′, reverse: 5’-ATAAGAATGCGGCCGCCTACAGGTTGCTGACGTAGGAC-3′ from pCav1.2 (addgene, Plasmid #26572), then the fragment was cloned into the vector pCav1.2 by restriction enzymes KpnI and NotI to construct a recombinant plasmid pCav1.2-KN. DsRed was amplified by primers: 5′-GGGAGACCCAAGCTGGCTAGC ATGGACAACACCGAGGACGTCAT3′, reverse: 5′-GTTTTCATTGACCATGGTACC CTGGGAGCCGGAGTGGCG from pLVX-EF1α-DsRed-monomer-N1 (Biovector NTCC Inc., Beijing, China). Finally, the DsRed fragment was ligased with linearized pCav1.2-KN by ExnaseII to construct the pDsRed-Cav1.2.

### Transfection

To transfect with small interfering RNA, BMSCs (0.5 × 10^6^) were seeded on a 6-well culture plate and transfected with Cav1.2 siRNA or Cav1.3 siRNA (Santa Cruz, Dallas, Texas, USA) using X-tremeGENE siRNA Transfection Reagent (Roche, Basel, Switzerland) according to the manufacturer’s instructions.

In order to down-regulate or overexpress ALPL in BMSCs, we firstly produced lentivirus, which respectively carried shALPL or ALPL cDNA. The Lentiviral vector and the ViraPower Packaging Mix were cotransfected into the 293T cell line to produce a lentiviral stock, following the protocol provided by the manufacturer. Virus-containing supernatants were harvested 48 h after transfection, pooled and filtered through 0.45-µm filters. Cells were treated with lentivirus at a multiplicity of infection of 100 at 37°C and 5% CO_2_. Plates were swirled every 15 min and fresh medium was added after 1 hour of incubation. The cells were exposed to lentivirus for 48 hours, followed by protein extraction for Western immunoblotting or differentiation induction.

For plasmid transfection of pCatenin-EGFP, pCav1.2 (addgene, Plasmid #26572), pCav1.3 (addgene, Plasmid #49332), DN-Dyn1 (addgene, Plasmid #55795), pCa_V_α2δ1(addgene, Plasmid #26575) and pCa_V_α2δ1mut (addgene, Plasmid #58730), we operated according to the protocol of X-tremeGENE HP DNA Transfection Reagent (Roche, Basel, Switzerland). After transfection, cells were used for protein extraction for western immunoblotting or for differentiation induction or for confocal images.

### MicroCT analysis

Femurs were harvested and analyzed using a desktop micro-CT system (eXplore Locus SP, GE Healthcare, USA). The scanner was set at a voltage of 80 kVp, a current of 80 μA and a resolution of 8 μm per pixel. Cross-sectional images of the femurmid-diaphysis were used to perform three-dimensional histomorphometric analysis of trabecular bone. Cross sectional volumetric bone mineral density (BMD) was measured for the right femur mid-diaphysis with a density phantom. Using 3-dimensional images, a region of interest in the secondary spongiosa was manually drawn near the endocortical surface. Bone volume relative to tissue volume (BV/TV) and trabecular bone number (Tb.N) were assessed as cancellous bone morphometric parameters.

### Histology

To assess the areas of trabecular bone and bone marrow, femurs and tibias were fixed in 4% paraformaldehyde (Sigma-Aldrich, St. Louis., MO, USA) and then decalcified with 5 % EDTA (pH 7.4), followed by paraffin embedding. The 6 μm paraffin sections were stained with hematoxylin and eosin (H&E) and analyzed using NIH ImageJ software. To label the mineralization fronts, the mice were given intraperitoneal injections of calcein (Sigma-Aldrich, USA, 20 mg/Kg body weight) at day 10 and day 3 before sacrifice. Bone dynamic histomorphometric analyses for MAR were performed according to the standardized nomenclature for bone histomorphometry under fluorescence microscopy (Leica DM 6000B, German).

### *In vivo* Oil Red O staining

To assess the adipose tissue in trabecular areas, femurs were fixed in 4% paraformaldehyde and decalcified with 5% EDTA (pH 7.4), followed by cryosectioning. Sections were stained with Oil Red O, and positive areas were quantified under microscopy and shown as a percentage of the total area. Briefly, sections were washed with 60% isopropanoland incubated in freshly made Oil Red O staining solution for 15min at room temperature and counterstained with hematoxylin. All regaents for Oil Red O staining were purchased from Sigma-Aldrich.

### Ionomycin treatment

Ionomycin (Alomone, Jerusalem, Israel) were dissolved in DMSO. For *in vivo* treatment, ionomycin was intraperitoneally administered to 12 weeks *alpl*^*+/-*^ mice and *alpl*^*-/-*^ CKO mice at a dose of 1mg/kg/day for 28 days. The control mice were treated with vehicle only. After ionomycin treatment, all groups of mice were healthy.

### Statistics

All experimental group sizes were chosen to ensure adequate statistical power despite the highly variable nature of the studies performed. No animals were excluded, and animals were randomly assigned groups for the studies. Experiments were not performed in a blinded fashion. Data were assessed for normal distribution and similar variance between groups. Comparisons between two groups were performed using independent unpaired two-tailed Student’s *t*-test, and the comparisons between more than two groups were analyzed using one-way ANOVA with the Bonferroni adjustment. *P* values less than 0.05 were considered statistically significant.

## Acknowledgements

This study was funded by the grants from National Natural Science Foundation of China (Nos. 81620108007 and 81870768), the National Key Research and Development Program of China (Nos.2016YFC1101400, Nos.2017YFA0104800), the Scientific Young Alma of Shaanxi province (2018KJXX-015) and Schoenleber pilot grant from University of Pennsylvania School of Dental Medicine.

## Author contributions

B.L., X.H. and Z.D. performed the animal experiments, immunofluorescence staining and immunoblot, cultured cells, collected data, analyzed the data and produced all figures and tables. K.X., collected the cells from patients, W.S. and L.G. helped with the immunofluorescence staining and participated in the animal surgery. S.L., W.L. C.H. and Y.Z. helped with data analysis. S.S. and Y.J. designed the experiments, oversaw the collection of results and data interpretation and drafted the reports. All authors have seen and approved the final version.

## References

Ambrosi TH, Scialdone A, Graja A, Gohlke S, Jank AM, Bocian C, Woelk L, Fan H, Logan DW, Schurmann A, Saraiva LR, Schulz TJ (2017) Adipocyte Accumulation in the Bone Marrow during Obesity and Aging Impairs Stem Cell-Based Hematopoietic and Bone Regeneration. Cell Stem Cell 20(6):771–784 e776

Barcia JP, Strife CF, Langman CB (1997) Infantile hypophosphatasia: treatment options to control hypercalcemia, hypercalciuria, and chronic bone demineralization. J Pediatr 130(5):825–828

Barradas AM, Fernandes HA, Groen N, Chai YC, Schrooten J, van de Peppel J, van Leeuwen JP, van Blitterswijk CA, de Boer J (2012) A calcium-induced signaling cascade leading to osteogenic differentiation of human bone marrow-derived mesenchymal stromal cells. Biomaterials 33(11):3205–3215

Belachew D, Kazmerski T, Libman I, Goldstein AC, Stevens ST, Deward S, Vockley J, Sperling MA, Balest AL (2013) Infantile hypophosphatasia secondary to a novel compound heterozygous mutation presenting with pyridoxine-responsive seizures. JIMD Rep 11:17–24

Berridge MJ, Lipp P, Bootman MD (2000) The versatility and universality of calcium signalling. Nat Rev Mol Cell Biol 1(1):11–21

Canti C, Nieto-Rostro M, Foucault I, Heblich F, Wratten J, Richards MW, Hendrich J, Douglas L, Page KM, Davies A, Dolphin AC (2005) The metal-ion-dependent adhesion site in the Von Willebrand factor-A domain of alpha2delta subunits is key to trafficking voltage-gated Ca2+ channels. Proc Natl Acad Sci U S A 102(32):11230– 11235

Catterall WA (2000) Structure and regulation of voltage-gated Ca2+ channels. Annu Rev Cell Dev Biol 16:521–555

Ciancaglini P, Yadav MC, Simao AM, Narisawa S, Pizauro JM, Farquharson C, Hoylaerts MF, Millan JL (2010) Kinetic analysis of substrate utilization by native and TNAP-, NPP1-, or PHOSPHO1-deficient matrix vesicles. J Bone Miner Res 25(4):716–723

Cui M, Li Q, Johnson R, Fleet JC (2012) Villin promoter-mediated transgenic expression of transient receptor potential cation channel, subfamily V, member 6 (TRPV6) increases intestinal calcium absorption in wild-type and vitamin D receptor knockout mice. J Bone Miner Res 27(10):2097–2107

Foster BL, Kuss P, Yadav MC, Kolli TN, Narisawa S, Lukashova L, Cory E, Sah RL, Somerman MJ, Millan JL (2017) Conditional Alpl Ablation Phenocopies Dental Defects of Hypophosphatasia. J Dent Res 96(1):81–91

Foster BL, Nagatomo KJ, Tso HW, Tran AB, Nociti FH, Jr., Narisawa S, Yadav MC, McKee MD, Millan JI, Somerman MJ (2013) Tooth root dentin mineralization defects in a mouse model of hypophosphatasia. J Bone Miner Res 28(2):271–282

Foster BL, Ramnitz MS, Gafni RI, Burke AB, Boyce AM, Lee JS, Wright JT, Akintoye SO, Somerman MJ, Collins MT (2014) Rare bone diseases and their dental, oral, and craniofacial manifestations. J Dent Res 93(7 Suppl): 7S–19S

Green EM, Barrett CF, Bultynck G, Shamah SM, Dolmetsch RE (2007) The tumor suppressor eIF3e mediates calcium-dependent internalization of the L-type calcium channel CaV1.2. Neuron 55(4):615–632

Hell JW, Westenbroek RE, Warner C, Ahlijanian MK, Prystay W, Gilbert MM, Snutch TP, Catterall WA (1993) Identification and differential subcellular localization of the neuronal class C and class D L-type calcium channel alpha 1 subunits. J Cell Biol 123(4):949–962

Hoenderop JG, van Leeuwen JP, van der Eerden BC, Kersten FF, van der Kemp AW, Merillat AM, Waarsing JH, Rossier BC, Vallon V, Hummler E, Bindels RJ (2003) Renal Ca2+ wasting, hyperabsorption, and reduced bone thickness in mice lacking TRPV5. J Clin Invest 112(12):1906–1914

J Ou, Sasaki H, Morimoto S, Kusakari Y, Shinji H, Obata T, Hongo K, Komukai K, Kurihara S (2008) Interaction of alpha1-adrenoceptor subtypes with different G proteins induces opposite effects on cardiac L-type Ca2+ channel. Circ Res 102(11):1378–1388

Jung H, Best M, Akkus O (2015) Microdamage induced calcium efflux from bone matrix activates intracellular calcium signaling in osteoblasts via L-type and T-type voltage-gated calcium channels. Bone 76:88–96

Li CJ, Xiao Y, Yang M, Su T, Sun X, Guo Q, Huang Y, Luo XH (2018) Long noncoding RNA Bmncr regulates mesenchymal stem cell fate during skeletal aging. J Clin Invest 128(12):5251–5266

Liu W, Zhang L, Xuan K, Hu C, Liu S, Liao L, Li B, Jin F, Shi S, Jin Y (2018) Alpl prevents bone ageing sensitivity by specifically regulating senescence and differentiation in mesenchymal stem cells. Bone Res 6:27

Liu Y, Yang R, Liu X, Zhou Y, Qu C, Kikuiri T, Wang S, Zandi E, Du J, Ambudkar IS, Shi S (2014) Hydrogen sulfide maintains mesenchymal stem cell function and bone homeostasis via regulation of Ca(2+) channel sulfhydration. Cell Stem Cell 15(1):66–78

Ludwig A, Flockerzi V, Hofmann F (1997) Regional expression and cellular localization of the alpha1 and beta subunit of high voltage-activated calcium channels in rat brain. J Neurosci 17(4):1339–1349

Merritt JE, McCarthy SA, Davies MP, Moores KE (1990) Use of fluo-3 to measure cytosolic Ca2+ in platelets and neutrophils. Loading cells with the dye, calibration of traces, measurements in the presence of plasma, and buffering of cytosolic Ca2+. Biochem J 269(2):513–519

Millan JL, Whyte MP (2016) Alkaline Phosphatase and Hypophosphatasia. Calcif Tissue Int 98(4):398–416

Moffat J, Grueneberg DA, Yang X, Kim SY, Kloepfer AM, Hinkle G, Piqani B, Eisenhaure TM, Luo B, Grenier JK, Carpenter AE, Foo SY, Stewart SA, Stockwell BR, Hacohen N, Hahn WC, Lander ES, Sabatini DM, Root DE (2006) A lentiviral RNAi library for human and mouse genes applied to an arrayed viral high-content screen. Cell 124(6):1283–1298

Narisawa S, Frohlander N, Millan JL (1997) Inactivation of two mouse alkaline phosphatase genes and establishment of a model of infantile hypophosphatasia. Dev Dyn 208(3):432–446

Praefcke GJ, McMahon HT (2004) The dynamin superfamily: universal membrane tubulation and fission molecules? Nat Rev Mol Cell Biol 5(2):133–147

Russell RG (1965) Excretion of Inorganic Pyrophosphate in Hypophosphatasia. Lancet 2(7410):461–464

Scheibe RJ, Kuehl H, Krautwald S, Meissner JD, Mueller WH (2000) Ecto-alkaline phosphatase activity identified at physiological pH range on intact P19 and HL-60 cells is induced by retinoic acid. J Cell Biochem 76(3):420–436

Seefried L, Baumann J, Hemsley S, Hofmann C, Kunstmann E, Kiese B, Huang Y, Chivers S, Valentin MA, Borah B, Roubenoff R, Junker U, Jakob F (2017) Efficacy of anti-sclerostin monoclonal antibody BPS804 in adult patients with hypophosphatasia. J Clin Invest 127(6):2148–2158

Snutch TP, Reiner PB (1992) Ca2+ channels: diversity of form and function. Curr Opin Neurobiol 2(3):247–253

Sun X, Kishore V, Fites K, Akkus O (2012) Osteoblasts detect pericellular calcium concentration increase via neomycin-sensitive voltage gated calcium channels. Bone 51(5):860–867

Takimoto K, Li D, Nerbonne JM, Levitan ES (1997) Distribution, splicing and glucocorticoid-induced expression of cardiac alpha 1C and alpha 1D voltage-gated Ca2+ channel mRNAs. J Mol Cell Cardiol 29(11):3035–3042

Tetreault MP, Bourdin B, Briot J, Segura E, Lesage S, Fiset C, Parent L (2016) Identification of Glycosylation Sites Essential for Surface Expression of the CaValpha2delta1 Subunit and Modulation of the Cardiac CaV1.2 Channel Activity. J Biol Chem 291(9):4826–4843

Tsien RW, Lipscombe D, Madison DV, Bley KR, Fox AP (1988) Multiple types of neuronal calcium channels and their selective modulation. Trends Neurosci 11(10):431–438

Waymire KG, Mahuren JD, Jaje JM, Guilarte TR, Coburn SP, MacGregor GR (1995) Mice lacking tissue non-specific alkaline phosphatase die from seizures due to defective metabolism of vitamin B-6. Nat Genet 11(1):45–51

Wen L, Wang Y, Wang H, Kong L, Zhang L, Chen X, Ding Y (2012) L-type calcium channels play a crucial role in the proliferation and osteogenic differentiation of bone marrow mesenchymal stem cells. Biochem Biophys Res Commun 424(3):439–445

Whyte MP (2016) Hypophosphatasia - aetiology, nosology, pathogenesis, diagnosis and treatment. Nat Rev Endocrinol 12(4):233–246

Whyte MP, Greenberg CR, Salman NJ, Bober MB, McAlister WH, Wenkert D, Van Sickle BJ, Simmons JH, Edgar TS, Bauer ML, Hamdan MA, Bishop N, Lutz RE, McGinn M, Craig S, Moore JN, Taylor JW, Cleveland RH, Cranley WR, Lim R, Thacher TD, Mayhew JE, Downs M, Millan JL, Skrinar AM, Crine P, Landy H (2012) Enzyme-replacement therapy in life-threatening hypophosphatasia. N Engl J Med 366(10):904–913

Whyte MP, Mahuren JD, Vrabel LA, Coburn SP (1985) Markedly increased circulating pyridoxal-5’-phosphate levels in hypophosphatasia. Alkaline phosphatase acts in vitamin B6 metabolism. J Clin Invest 76(2):752–756

Whyte MP, Rockman-Greenberg C, Ozono K, Riese R, Moseley S, Melian A, Thompson DD, Bishop N, Hofmann C (2016) Asfotase Alfa Treatment Improves Survival for Perinatal and Infantile Hypophosphatasia. J Clin Endocrinol Metab 101(1):334–342

Zhang J, Li M, Kang ET, Neoh KG (2016) Electrical stimulation of adipose-derived mesenchymal stem cells in conductive scaffolds and the roles of voltage-gated ion channels. Acta Biomater 32:46–56

Zimmermann H (2006) Nucleotide signaling in nervous system development. Pflugers Arch 452(5):573–588

McLaughlin RN, Poelwijk FJ, Raman A, Gosal WS, Ranganathan R. The spatial architecture of protein function and adaptation. Nature. 2012 Nov;491(7422):138–142.

Firnberg E, Labonte JW, Gray JJ, Ostermeier M. A comprehensive, high-resolution map of a gene’s fitness landscape. Mol Biol Evol. 2014 Jun;31(6):1581–1592.

Boucher JI, Bolon DN, Tawfik DS. Quantifying and understanding the fitness effects of protein mutations: Laboratory versus nature. Protein Sci. 2016 07;25(7):1219–1226.

Hopf TA, Ingraham JB, Poelwijk FJ, Schärfe CPI, Springer M, Sander C, et al. Mutation effects predicted from sequence co-variation. Nature Biotechnology. 2017 01;35:128 EP–.

Figliuzzi M, Jacquier H, Schug A, Tenaillon O, Weigt M. Coevolutionary Landscape Inference and the Context-Dependence of Mutations in Beta-Lactamase TEM-1. Mol Biol Evol. 2016 Jan;33(1):268–280.

Mann JK, Barton JP, Ferguson AL, Omarjee S, Walker BD, Chakraborty A, et al. The fitness landscape of HIV-1 gag: advanced modeling approaches and validation of model predictions by in vitro testing. PLoS Comput Biol. 2014 Aug;10(8):e1003776.

Sim NL, Kumar P, Hu J, Henikoff S, Schneider G, Ng PC. SIFT web server: predicting effects of amino acid substitutions on proteins. Nucleic Acids Res. 2012 Jul;40(Web Server issue):W452–457.

Dehouck Y, Kwasigroch JM, Gilis D, Rooman M. PoPMuSiC 2.1: a web server for the estimation of protein stability changes upon mutation and sequence optimality. BMC Bioinformatics. 2011 May;12:151.

Adzhubei IA, Schmidt S, Peshkin L, Ramensky VE, Gerasimova A, Bork P, et al. A method and server for predicting damaging missense mutations. Nat Methods. 2010 Apr;7(4):248–249.

Cheng J, Randall A, Baldi P. Prediction of protein stability changes for single-site mutations using support vector machines. Proteins. 2006 Mar;62(4):1125–1132.

Capriotti E, Fariselli P, Casadio R. I-Mutant2.0: predicting stability changes upon mutation from the protein sequence or structure. Nucleic Acids Res. 2005 Jul;33(Web Server issue):W306–310.

Ng PC, Henikoff S. SIFT: Predicting amino acid changes that affect protein function. Nucleic Acids Res. 2003 Jul;31(13):3812–3814.

Breen MS, Kemena C, Vlasov PK, Notredame C, Kondrashov FA. Epistasis as the primary factor in molecular evolution. Nature. 2012 Oct;490(7421):535–538.

McCandlish DM, Shah P, Plotkin JB. Epistasis and the Dynamics of Reversion in Molecular Evolution. Genetics. 2016 07;203(3):1335–1351.

Stein RR, Marks DS, Sander C. Inferring Pairwise Interactions from Biological Data Using Maximum-Entropy Probability Models. PLoS Comput Biol. 2015 Jul;11(7):e1004182.

Weigt M, White RA, Szurmant H, Hoch JA, Hwa T. Identification of direct residue contacts in protein-protein interaction by message passing. Proc Natl Acad Sci USA. 2009 Jan;106(1):67–72.

Mihalek I, Res I, Lichtarge O. A family of evolution-entropy hybrid methods for ranking protein residues by importance. J Mol Biol. 2004;336(5):1265–1282.

Lichtarge O, Bourne HR, Cohen FE. An evolutionary trace method defines binding surfaces common to protein families. J Mol Biol. 1996;257(2):342–358.

Engelen S, Trojan LA, Sacquin-Mora S, Lavery R, Carbone A. Joint evolutionary trees: a large-scale method to predict protein interfaces based on sequence sampling. PLoS Comput Biol. 2009;5(1):e1000267.

Laine E, Carbone A. Local Geometry and Evolutionary Conservation of Protein Surfaces Reveal the Multiple Recognition Patches in Protein-Protein Interactions. PLoS Comput Biol. 2015 Dec;11(12):e1004580.

Ripoche H, Laine E, Ceres N, Carbone A. JET2 Viewer: a database of predicted multiple, possibly overlapping, protein-protein interaction sites for PDB structures. Nucleic Acids Res. 2017 Jan;45(D1):D236–D242.

Karami Y, Bitard-Feildel T, Laine E, Carbone A. “Infostery”analysis of short molecular dynamics simulations identifies highly sensitive residues and predicts deleterious mutations. Scientific Reports. 2018;8(1):16126.

Neher RA, Bedford T. Real-Time Analysis and Visualization of Pathogen Sequence Data. J Clin Microbiol. 2018 Nov;56(11).

Altschul SF, Madden TL, Schäffer AA, Zhang J, Zhang Z, Miller W, et al. Gapped BLAST and PSI-BLAST: a new generation of protein database search programs. Nucleic Acids Research. 1997 09;25(17):3389–3402.

Studier JA, Keppler KJ. A note on the neighbor-joining algorithm of Saitou and Nei. Mol Biol Evol. 1988 Nov;5(6):729–731.

Laine E, Carbone A. The geometry of protein-protein interfaces reveals the multiple origins of recognition patches. PLoS Computational Biology. 2015;11(12):e1004580.

Edgar RC. MUSCLE: multiple sequence alignment with high accuracy and high throughput. Nucleic Acids Res. 2004;32(5):1792–1797.

Romero PA, Tran TM, Abate AR. Dissecting enzyme function with microfluidic-based deep mutational scanning. Proc Natl Acad Sci USA. 2015 Jun;112(23):7159–7164.

Roscoe BP, Bolon DN. Systematic exploration of ubiquitin sequence, E1 activation efficiency, and experimental fitness in yeast. J Mol Biol. 2014 Jul;426(15):2854–2870.

Roscoe BP, Thayer KM, Zeldovich KB, Fushman D, Bolon DN. Analyses of the effects of all ubiquitin point mutants on yeast growth rate. J Mol Biol. 2013 Apr;425(8):1363–1377.

Melamed D, Young DL, Gamble CE, Miller CR, Fields S. Deep mutational scanning of an RRM domain of the Saccharomyces cerevisiae poly(A)-binding protein. RNA. 2013 Nov;19(11):1537–1551.

Stiffler MA, Hekstra DR, Ranganathan R. Evolvability as a function of purifying selection in TEM-1 Îš-lactamase. Cell. 2015 Feb;160(5):882–892.

Kitzman JO, Starita LM, Lo RS, Fields S, Shendure J. Massively parallel single-amino-acid mutagenesis. Nat Methods. 2015 Mar;12(3):203–206.

Melnikov A, Rogov P, Wang L, Gnirke A, Mikkelsen TS. Comprehensive mutational scanning of a kinase in vivo reveals substrate-dependent fitness landscapes. Nucleic Acids Res. 2014 Aug;42(14):e112.

Araya CL, Fowler DM, Chen W, Muniez I, Kelly JW, Fields S. A fundamental protein property, thermodynamic stability, revealed solely from large-scale measurements of protein function. Proc Natl Acad Sci USA. 2012 Oct;109(42):16858–16863.

Firnberg E, Labonte JW, Gray JJ, Ostermeier M. A Comprehensive, High-Resolution Map of a Gene’s Fitness Landscape. Mol Biol Evol. 2016 05;33(5):1378.

Rockah-Shmuel L, Toth-Petroczy A, Tawfik DS. Systematic Mapping of Protein Mutational Space by Prolonged Drift Reveals the Deleterious Effects of Seemingly Neutral Mutations. PLoS Comput Biol. 2015 Aug;11(8):e1004421.

Jacquier H, Birgy A, Le Nagard H, Mechulam Y, Schmitt E, Glodt J, et al. Capturing the mutational landscape of the beta-lactamase TEM-1. Proc Natl Acad Sci USA. 2013 Aug;110(32):13067–13072.

Qi H, Olson CA, Wu NC, Ke R, Loverdo C, Chu V, et al. A quantitative high-resolution genetic profile rapidly identifies sequence determinants of hepatitis C viral fitness and drug sensitivity. PLoS Pathog. 2014 Apr;10(4):e1004064.

Wu NC, Olson CA, Du Y, Le S, Tran K, Remenyi R, et al. Functional Constraint Profiling of a Viral Protein Reveals Discordance of Evolutionary Conservation and Functionality. PLoS Genet. 2015 Jul;11(7):e1005310.

Mishra P, Flynn JM, Starr TN, Bolon DNA. Systematic Mutant Analyses Elucidate General and Client-Specific Aspects of Hsp90 Function. Cell Rep. 2016 Apr;15(3):588–598.

Deng Z, Huang W, Bakkalbasi E, Brown NG, Adamski CJ, Rice K, et al. Deep sequencing of systematic combinatorial libraries reveals Îš-lactamase sequence constraints at high resolution. J Mol Biol. 2012 Dec;424(3-4):150–167.

Starita LM, Pruneda JN, Lo RS, Fowler DM, Kim HJ, Hiatt JB, et al. Activity-enhancing mutations in an E3 ubiquitin ligase identified by high-throughput mutagenesis. Proc Natl Acad Sci USA. 2013 Apr;110(14):E1263–1272.

Aakre CD, Herrou J, Phung TN, Perchuk BS, Crosson S, Laub MT. Evolving new protein-protein interaction specificity through promiscuous intermediates. Cell. 2015 Oct;163(3):594–606.

Doud MB, Bloom JD. Accurate Measurement of the Effects of All Amino-Acid Mutations on Influenza Hemagglutinin. Viruses. 2016 06;8(6).

Peterson EL, Kondev J, Theriot JA, Phillips R. Reduced amino acid alphabets exhibit an improved sensitivity and selectivity in fold assignment. Bioinformatics. 2009;25(11):1356–1362. Available from: +http://dx.doi.org/10.1093/bioinformatics/btp164.

Andersen CAF, Brunak S. Representation of Protein-sequence Information by Amino Acid Subalphabets. AI Mag. 2004 Mar;25(1):97–104. Available from: http://dl.acm.org/citation.cfm?id=996917.996927.

Miyazawa S, Jernigan RL. Residue-residue potentials with a favorable contact pair term and an unfavorable high packing density term, for simulation and threading. J Mol Biol. 1996 Mar;256(3):623–644.

Cieplak M, Holter NS, Maritan A, Banavar JR. Amino acid classes and the protein folding problem. The Journal of Chemical Physics. 2001;114(3):1420–1423. Available from: https://doi.org/10.1063/1.1333025.

Solis AD, Rackovsky S. Optimized representations and maximal information in proteins. Proteins. 2000 Feb;38(2):149–164.

Prlic A, Domingues FS, Sippl MJ. Structure-derived substitution matrices for alignment of distantly related sequences. Protein Eng. 2000 Aug;13(8):545–550.

Landes C, Risler JL. Fast databank searching with a reduced amino-acid alphabet. Comput Appl Biosci. 1994 Jul;10(4):453–454.

Li T, Fan K, Wang J, Wang W. Reduction of protein sequence complexity by residue grouping. Protein Eng. 2003 May;16(5):323–330.

Liu X, Liu D, Qi J, Zheng WM. Simplified amino acid alphabets based on deviation of conditional probability from random background. Phys Rev E. 2002 Aug;66:021906. Available from: https://link.aps.org/doi/10.1103/PhysRevE.66.021906.

Murphy LR, Wallqvist A, Levy RM. Simplified amino acid alphabets for protein fold recognition and implications for folding. Protein Eng. 2000 Mar;13(3):149–152.

Johnson MS, Overington JP. A structural basis for sequence comparisons. An evaluation of scoring methodologies. J Mol Biol. 1993 Oct;233(4):716–738.

Melo F, Marti-Renom MA. Accuracy of sequence alignment and fold assessment using reduced amino acid alphabets. Proteins. 2006 Jun;63(4):986–995.

Mirny LA, Shakhnovich EI. Universally conserved positions in protein folds: reading evolutionary signals about stability, folding kinetics and function. J Mol Biol. 1999 Aug;291(1):177–196.

Thomas PD, Dill KA. An iterative method for extracting energy-like quantities from protein structures. Proc Natl Acad Sci USA. 1996 Oct;93(21):11628–11633.

Wang J, Wang W. A computational approach to simplifying the protein folding alphabet. Nat Struct Biol. 1999 Nov;6(11):1033–1038.

Henikoff S, Henikoff JG. Amino acid substitution matrices from protein blocks. Proc Natl Acad Sci USA. 1992 Nov;89(22):10915–10919.

